# The transcription factor Zeb2 drives formation of age-associated B cells

**DOI:** 10.1101/2021.07.24.453633

**Authors:** Dai Dai, Shuangshuang Gu, Xiaxia Han, Huihua Ding, Yang Jiang, Xiaoou Zhang, Chao Yao, Soonmin Hong, Jinsong Zhang, Yiwei Shen, Guojun Hou, Bo Qu, Haibo Zhou, Yuting Qin, Yuke He, Jianyang Ma, Zhihua Yin, Zhizhong Ye, Jie Qian, Qian Jiang, Lihua Wu, Qiang Guo, Sheng Chen, Chuanxin Huang, Leah C. Kottyan, Matthew T. Weirauch, Carola G. Vinuesa, Nan Shen

**Author notes:** Corresponding author. Email: Nan Shen Carola G. Vinuesa. These authors contributed equally to the work (Dai Dai, Shuangshuang Gu, Xiaxia Han).

## Abstract

Age-associated B-cells (ABCs) accumulate during infection, aging and autoimmunity, contributing to lupus pathogenesis. Here, we screen for transcription factors driving ABC formation and find Zeb2 is required for human and mouse ABC differentiation in-vitro. ABCs are reduced in ZEB2 haploinsufficient individuals and in mice lacking Zeb2 in B-cells. In mice with TLR7-driven lupus, Zeb2 is essential for ABC formation and autoimmune pathology. Zeb2 binds to the +20kb intronic enhancer of Mef2b, repressing Mef2b-mediated germinal center B-cell differentiation and promoting ABC formation. Zeb2 also targets genes important for ABC specification and function including *Itgax*. Zeb2-driven ABC differentiation requires Jak-Stat signaling, and treatment with the Jak1/3 inhibitor tofacitinib reduces ABC accumulation in autoimmune mice and patients. Zeb2 thus emerges as a driver of B-cell autoimmunity.

**One-Sentence Summary:** Zeb2 is essential for Age-associated B cells differentiation and function.

## Main Text

Age-associated B cells (ABC) characterized by high expression of CD11c and T-bet are found at increased numbers in aged female mice, infection models, and multiple autoimmune diseases (*1*). This novel B cell subset has been given various names including ABCs, CD11c^+^ B cells, CD11c^+^T-bet^+^ B cells, CD21^-^CD23^-^ B cells, Be1 cells, double negative B cells (DN), DN2 or atypical memory B cells (*2-10*). ABCs or ABC-like B cells identified as CD11c^+^T-bet^+^CD11b^+^CD21^-^ CD23^-^CD86^hi^MHCII^hi^CD95^+^ in mice (*2, 8*) and CD11c^+^T-bet^+^CD27^-^IgD^-^CD38^-^CD21^-^CD95^hi^ FCRL5^+^ in humans (*4, 6*) will be referred to as ABCs henceforth, following the suggestion of a recent review (*1*). In autoimmune settings, these B cells are enriched for autoantibody specificities. It is widely accepted that these cells are antigen-experienced and it has been suggested that they can persist in tissues and rapidly differentiate into antibody-secreting cells (ASCs) upon antigen reencounter or innate stimulation (*11, 12*).

The transcription factors T-bet, IRF5 and IRF8 are highly expressed in ABCs and have been put forward as functional regulators of ABC differentiation (*5, 9, 10, 13*). However, except for IgG2a/c isotype switching, T-bet is dispensable for ABC accumulation or maintenance of ABC features (*13-15*), arguing against it being the master regulator of ABCs. IRF5 and IRF8 are broadly expressed in other B cell subsets and have been reported to be involved in cell activation, proliferation, differentiation, and function (*16-18*). In search for the transcription factor (TF) essential for ABC formation, we screened all TFs expressed by these cells and identified Zeb2 as the key regulator required for ABC specification and differentiation in mouse and human.

## Results

### Differentially-expressed TFs in autoimmune ABCs

To gain insights into the nature of ABCs, we sorted peripheral B cells from a patient with new-onset SLE (table S1) and performed droplet-based scRNA-seq. Seven distinct clusters containing from 94 to 1640 cells were revealed by unsupervised clustering with a two-dimensional uniform manifold approximation and projection (UMAP) (Fig. 1A and fig. S1, A and B). These clusters were assigned to known peripheral B cell subsets including transitional B cells (TrB), naïve B cells (NavB), activated naïve B cells (aNavB), ABCs, memory B cells (MemB), plasmablasts (PB), and plasma cells (PC) by comparing differentially-expressed genes with established markers and previously described gene expression profiles (*6, 19, 20*). Landmark genes were used to identify clusters (fig. S1, A and C). ABCs preferentially expressed genes encoding key surface markers *ITGAX, CD19, MS4A1, CD86, FCRLA, FCRL3/5*, and *FCGR2B*, and they lacked *CR2, FCER2, CD27, CXCR5* and *IGHD* (Fig. 1B and fig. S1C). Besides these, we identified 43 differentially-expressed TFs in ABCs: 27 upregulated and 16 downregulated (Fig. 1C).

**Figure 1.**
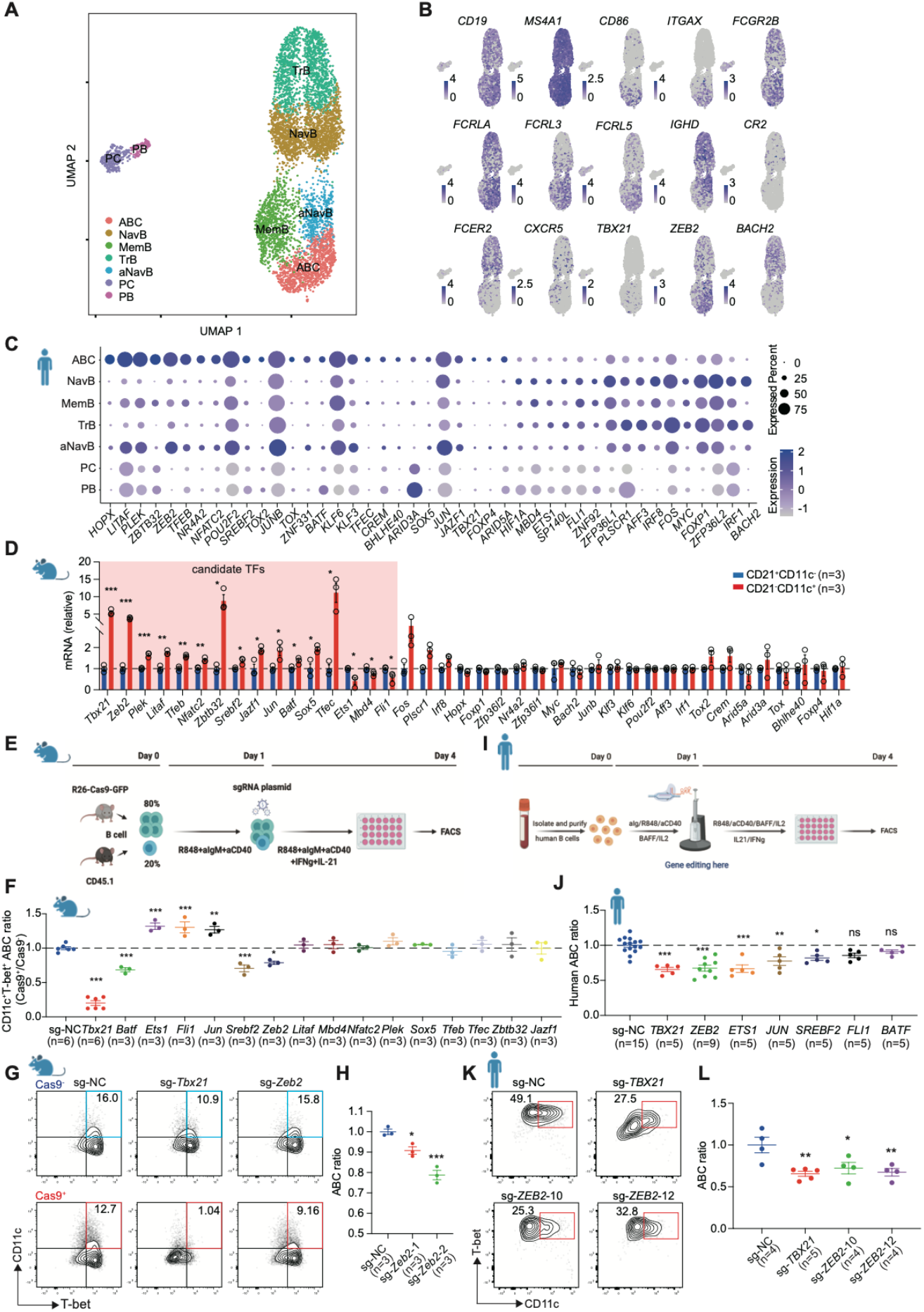
CRISPR/Cas9 based screen of transcription factors for ABC differentiation. **(A)** CD19^+^ B cells were sorted from PBMCs of new-onset SLE patient and performed droplet-based scRNA-seq. Seven clusters are defined as transitional B cell (TrB), naïve B cell (NavB), activate naïve B cell (aNavB), ABC, memory B cell (MemB), plasma blast (PB), and plasma cells (PC). Each point is a single cell colored by cluster assessment. (**B**) UMAP plots of select genes expression distinguishing ABCs. Color scales for each gene show log-normalized expression levels. (**C**) Dot plots of 43 differentially-expressed transcription factors in ABCs with 27 upregulated TFs and 16 downregulated TFs. Dot size represents the proportion of cells expressing indicated TF in particular cluster and intensity of color represents the average expression level. (**D**) C57BL/6J mice were intraperitoneally injected with CD4^+^ T cells from bm12 mice and induced for 2 weeks. Splenic CD19^+^CD11c^+^CD21^-^ and CD19^+^CD11c^-^CD21^+^ B cells were sorted by flow cytometry to validate mouse genes encoding equivalent TFs in (**C**). (**E**) Flow chart of Cas9 based screen and in vitro mouse ABC differentiation assay. On day 0, splenic B cells from Cas9-EGFP and B6-CD45.1 mouse were mixed and activated. On day 1, activated B cells were retroviral infected with sgRNA-plasmids and further cultured with ABC-skewing cocktail for 3 days. On day 4, cells were analyzed by flow cytometry. (**F**) Statistical analysis of ABC (CD11c^+^T-bet^+^) ratio in groups targeting indicated genes. (**G**) Flow cytometry plots of ABC (CD11c^+^T-bet^+^) ratio in retroviral infected B cells of sgRNA targeting *Tbx21* or *Zeb2*. (**H**) Statistical analysis of ABC (CD11c^+^T-bet^+^) ratio in retroviral infected B cells of two distinct sgRNAs targeting *Zeb2*. **(I)** Flow chart of Cas9-RNP based screen and in vitro human ABC differentiation assay. On day 0, human B cells were activated. On day 1, activated B cells was electronical delivered of Cas9-RNP and further cultured with ABC-skewing cocktail. On day 4, cells were analyzed by flow cytometry. (**J**) Statistical analysis of human ABCs (CD27^-^ IgD^-^CD11c^+^T-bet^+^) after gene editing and in vitro differentiation. ABC ratio was normalized with sg-NC. (**K-L**) Representative plots of human ABCs (CD27^-^IgD^-^CD11c^+^T-bet^+^) and statistical analysis of ABC ratio in sg-*TBX21* and two distinct sg-*ZEB2* edited human B cells comparing with sg-NC. N represents distinct samples (biological repeats). Data are representative of 3-4 independent experiments. Data are mean ± SEM values. *p<0.05, **P<0.01, ***P<0.001, ns, not significant, unpaired t-test (**D**) and ordinary one-way ANOVA with Dunnett’s multiple comparisons testing (**F, J, H** and **L**).

We previously reported that T-bet^+^CD11c^+^ B cells accumulate in bm12-induced lupus mice and contribute to autoantibody production (*3*). Flow cytometric staining revealed that CD11c^+^ B cells from bm12-induced mice exhibited ABC-specific features including a hyperactivated state (CD19^hi^CD86^hi^CD62L^low^CD44^hi^CD95^hi^MHCII^hi^), characteristic markers (CD11c^+^T-bet^hi^CD11b^hi^), specific lineage association (CD23^-^CD21^-^CD93^-^CD5^int^CD138^-^IgD^low^IgM^int^) and elevated expression of chemokine receptors CXCR3 and CXCR4 (fig. S2A). CD21 expression excludes the small fraction of naive CD11c^+^ B cells (fig. S2B). Consistent with previous reports, bm12-indued CD11c^+^CD21^-^ B cells expressed high amounts of *Cxcl9* and *Cxcl10* transcripts (Fig. S2C).

We then used qPCR to validate 40 differentially-expressed TFs identified in our human scRNA-seq data excluding *ZNF331, SP140L*, and *ZNF92*, which lack orthologs in mice (Fig.1D). Amongst these, 13 upregulated TFs (*Tbx21, Zeb2, Plek, Litaf, Tfeb, Nfatc2, Zbtb32, Srebf2, Jazf1, Jun, Sox5, Tfec, Batf*) and 3 downregulated TFs (*Ets1, Mbd4, Fli1*) were identically regulated in human and mouse ABCs and were therefore considered potential transcriptional regulators of ABC differentiation.

### Screen for TFs directing ABC differentiation

To screen for functional transcriptional regulators of ABC differentiation, B cells from Cas9 transgenic mice and congenic CD45.1 mice were cocultured in a 4:1 ratio and activated with anti-IgM, anti-CD40, and R848 for 24 hours. B cells expressing CD45.1 lacked Cas9 and served as controls in these co-cultures. Activated B cells were then retrovirally infected with the sgRNA plasmids co-expressing blue fluorescent protein (BFP) and cultured for a further 3 days with the ABC differentiation cocktail (*10, 21*) (Fig. 1E). To validate our genome editing tools we introduced a sgRNA sequence specific for *Itgax* and compared the ratio between live BFP^+^ CD45.1^-^ (Cas9^+^, edited) and BFP^+^ CD45.1^+^ (Cas9^-^, unedited) CD11c^+^ T-bet^+^ ABCs (fig. S3A). We found effective reduction of CD11c^+^T-bet^+^ ABC formation from Cas9^+^ B cells (fig. S3B).

We then screened the 16 candidate TFs and identified 7 (*T-bet, Batf, Ets1, Fli1, Jun, Srebf2, Zeb2*) that could significantly (P<0.05) alter ABC formation (Fig. 1, F and G, and fig. S3C). Except for *Zeb2*, ablation of the other 6 TFs predominantly influenced cell viability (fig. S3, D and E). Two different sgRNAs targeting *Zeb2* in separate Cas9^+^ B cell cultures led to reduced ABC formation (Fig. 1H), which excluded an off-target editing effect. Taken together, we identified 7 functional TFs influencing mouse ABC differentiation, with one of them, *Zeb2*, acting by means other than promoting cell survival.

Next, we set out to determine whether besides *Zeb2*, any of the other 6 TFs was required for human ABC lineage specification, using an established in vitro ABC induction protocol used in human studies (*4, 6, 9*). We transduced Cas9-guide RNA ribonucleoprotein (RNP) complexes by electroporation to edit genes in primary human B cells. Using a sgRNA sequence targeting *CD19* as a positive control, we achieved extremely efficient gene disruption (fig. S4A). Primary B cells were isolated and purified from the blood of healthy individuals. After activation for one day, cells were electroporated with RNP and treated with the ABC differentiation cocktail for another 3 days (Fig. 1I). Except for *BATF* and *FLI1*, all the other 5 TFs identified in mice led to changes in human ABCs when individually ablated (P<0.05) (Fig. 1, J and K, and fig. S4B). Two different guide RNAs targeting *ZEB2* achieved impaired ABC induction (Fig. 1L). Gene editing of *ETS1* and *JUN* had opposite effects on human ABCs and these TFs were excluded. Ablation of *TBX21, ZEB2*, and *SREBF2* dampened ABC differentiation in both human and mouse B cells. However, *TBX21* and *SREBF2* deficiency also led to altered B cell viability during in vitro induction. (fig. S4, C and D). As a key regulator of lipid metabolism, *SREBF2* is required to supply lipids for membrane synthesis and has been reported to influence T cell survival and the GC response (*22, 23*). Although there is no published evidence that T-bet controls B cell proliferation or death, T-bet has been implicated in B lymphoproliferative disorders (*24*). While it is possible that these genes may play additional roles beyond promoting cell viability, *Zeb2* was the single TF with consistent regulatory effects in human and mouse ABC formation unrelated to cell survival, and therefore a promising putative ABC transcriptional regulator.

### ZEB2 happloinsufficiency impairs human ABC formation

*Zeb2* belongs to the zinc-finger E homeobox-binding protein family and is known to drive epithelial to mesenchymal transition in early fetal development and cancer progression (*25*). Mutation of this gene is associated with Mowat-Wilson syndrome (MWS) (OMIM # 235730), a rare genetic disorder characterized by intellectual disability, distinctive facial features, seizures, and the predisposition to a gastrointestinal disease known as Hirschsprung disease (*26-28*). Over 100 heterozygous variants in the *ZEB2* gene have been reported in MWS patients, which are usually whole or partial gene deletions or truncating mutations, underscoring that haploinsufficiency is the main pathological mechanism for MWS (*29*). Even though *Zeb2* has been reported to be expressed in immune cells and participate in transcriptional networks involved in cell differentiation, maintenance, and function from mice studies (*30-38*), little is known about the immunological alterations caused by decreased function of ZEB2 in MWS patients.

We obtained PBMCs from 5 unrelated patients who had been diagnosed with MWS by the detection of *ZEB2* mutations (table S4) and their clinical manifestations (table S5). We performed whole-exome sequencing of these 5 families and confirmed *de novo* heterozygous germline *ZEB2* mutations in patient A (c.1027C>T, p.Arg343X), B (c.2851C>T, p.Gln951X), C (c.1005delT, p.Ile335Metfs*2), D (chr2:138434153-145285163 del) and E (chr2:145147017-145274917 del) (Fig. 2, A and B). Immunological investigations were performed on peripheral blood mononuclear cell samples for MWS patients and age/gender-matched healthy donors (HD). High-dimensional analysis of lymphocyte subsets by flow cytometry revealed the frequency of ABCs was dramatically decreased by *ZEB2* deficiency (Fig. 2C and fig. S5A). To further explore the ABC phenotype, B cell profiling was conducted in all MWS patients using an identical staining panel. Seven major B cell clusters were visually apparent and assigned according to the expression of the known markers (Fig. 2, D and E). The alterations of B cell subsets were compared between MWS patients and HDs (Fig. 2F and fig. S5B). Of note, the frequency of ABCs was significantly decreased in MWS patients by automated clustering (Fig. 2F) or manual gating with specific markers (Fig. 2G and fig. S5C). The frequency of activated naïve B cells, known as the ABC progenitors, was also significantly decreased (fig. S5, B and D) (*6*). The frequency of switched memory B cells decreased in MWS patient but not as dramatically as ABCs (fig. S5B). By contrast, DN1 B cells, which represent an alternative trajectory for B cell terminal differentiation, were increased in MWS patients (fig. S5B) (*6*). To further explore the regulatory effects of *Zeb2* on ABC formation, we isolated B cells from a patient with MWS and induced ABC differentiation in vitro. Consistent with the in vivo data, ABC formation was also impaired (Fig. 2H). This evidence from patients with *ZEB2* loss-of-function mutations confirms that this TF is required for human ABC formation both in vitro and in vivo.

**Figure 2.**
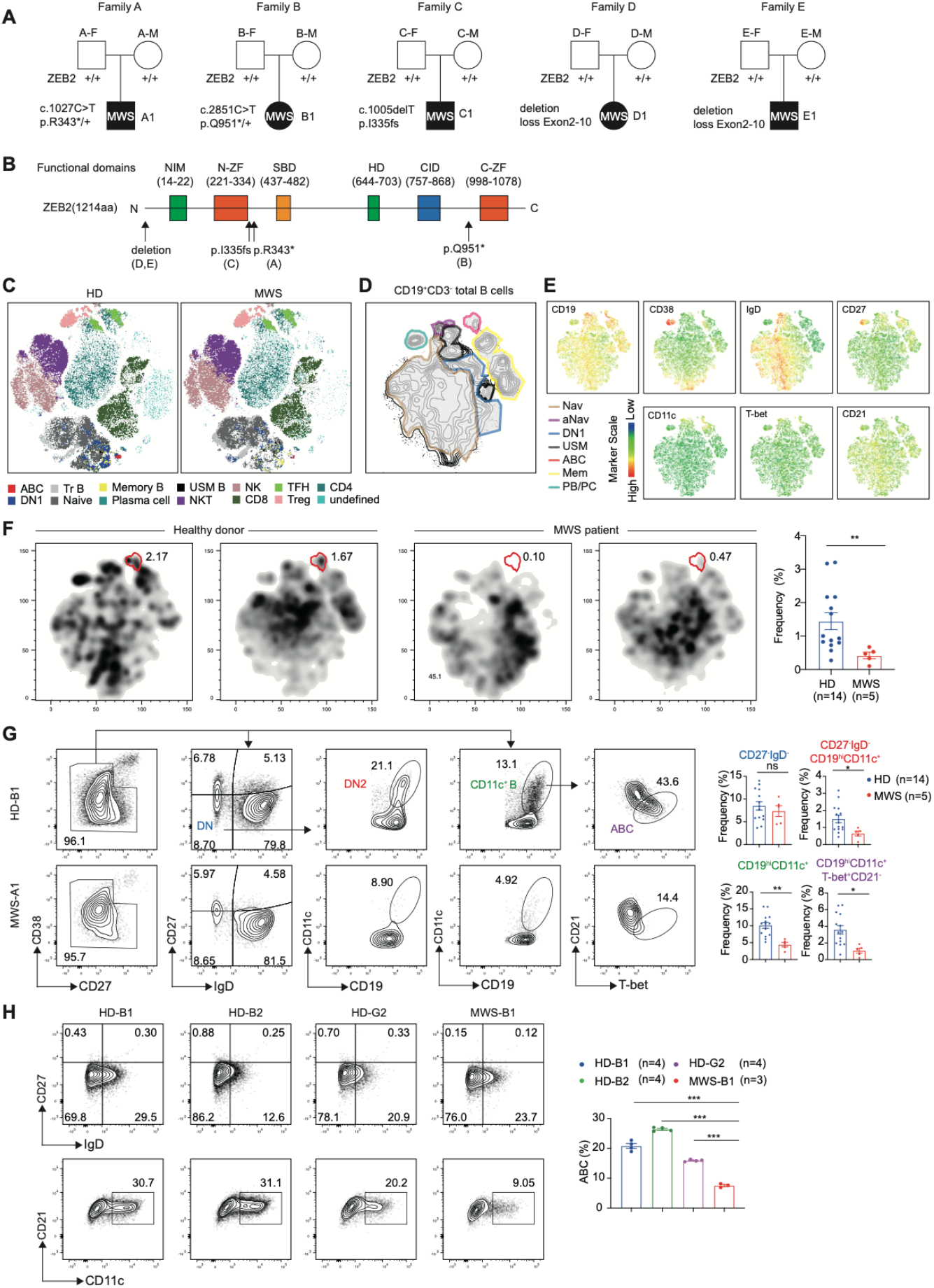
ZEB2 is required for human ABC formation. (**A**) Family pedigrees showing de novo heterozygous mutations of ZEB2 in Mowat-Wilson syndrome (MWS) patients. Family A shows ZEB2 nonsense mutation c.1027C>T (p. Arg343*), Family B shows ZEB2 nonsense mutation c.2851C>T (p. Gln951*), Family C shows ZEB2 frameshift variant c.1005delT (p. I335fs), Family D and E shows ZEB2 loss exon 2-10. (**B**) Schematic of the general, linear structure of the functional domain composition of ZEB2 protein. The black arrows show the location of ZEB2 mutation described in (**A**). (**C**) t-SNE plots of lymphocytes clusters in PBMC of healthy donor (HD) and MWS patient analyzed by flow cytometry. (**D**) Comprised t-SNE plots of peripheral B cell clusters for all donors. Seven B cell clusters were identified based on lineage marker expression as naïve B cell (Nav), activate naïve B cell (aNav), CXCR5^+^ double negative B cells (DN1), unswitch memory B cells (USM), age-associated B cells (ABC), memory B cell (Mem), plasma blast (PB) and plasma cells (PC). (**E**) t-SNE plots of peripheral B cells displaying CD19, CD38, IgD, CD27, CD11c, T-bet, and CD21 expression. (**F**) Separated t-SNE plots for two representative HD (left) and MWS (right) samples. Statistical analysis of ABC in PBMC for HD versus MWS group. (**G**) Representative plots and frequency of CD19^+^CD38^-^CD27^-^IgD^-^(DN), CD19^+^CD38^-^CD27^-^ IgD^-^CD11c^+^(DN2), CD19^+^CD38^low/mid^CD27^low/mid^CD11c^+^(CD11c^+^B) and CD19^+^CD38^low/mid^CD27^low/mid^IgD^-^ CD11c^+^CD21^-^T-bet^+^(ABC) in PBMC for HD versus MWS group. (**H**) Primary B cells were isolated from HDs and a MWS patient and cultured with ABC-skewing cocktail for 3 days. Representative plots and frequency of ABCs (CD19^+^CD38^-^CD27^-^IgD^-^CD11c^+^CD21^-^). n represents distinct samples (biological repeats except **H**). Data are mean ± SEM values. *p<0.05, **P<0.01, ***P<0.001, ns, not significant, unpaired t-test (**G, H**) with Welch’s correction (**F**).

### Zeb2 determines the pathogenic roles of ABCs in lupus autoimmunity

To further explore how *Zeb2* deficiency affects ABC formation we generated mice selectively lacking *Zeb2* in B cells by crossing *Zeb2* floxed mice with CD19-cre mice (fig. S6, A to C). We first isolated B cells from Zeb2^f/f^CD19^cre/+^ B cell conditional knockout mice (B-Zeb2^KO^), Zeb2^f/+^ CD19^cre/+^ heterozygous mice (B-Zeb2^Het^), Zeb2^f/f^ CD19^+/+^ control mice (floxed-Ctrl) and Zeb2^+/+^ CD19^cre/+^ (CD19-Ctrl) and cultured them for 3 days in ABC-skewing conditions. *Zeb2* deficiency in B cells reduced ABC differentiation by over 50% (Fig. 3A). Even mice hemizygous for *Zeb2* in B cells displayed a reduction in ABC formation, which was not as severe as that in B-Zeb2^KO^ B cells. Next, we investigated whether retroviral overexpression of *Zeb2* in B cells could promote ABC development. Splenic B cells were transduced with plasmids expressing *Zeb2* or *Tbx21*, and then cultured in ABC-skewing conditions. Both *Zeb2* and *Tbx21* promoted ABC formation, indicating that either of these transcription factors is sufficient for ABC differentiation in vitro (Fig. 3B).

**Figure 3.**
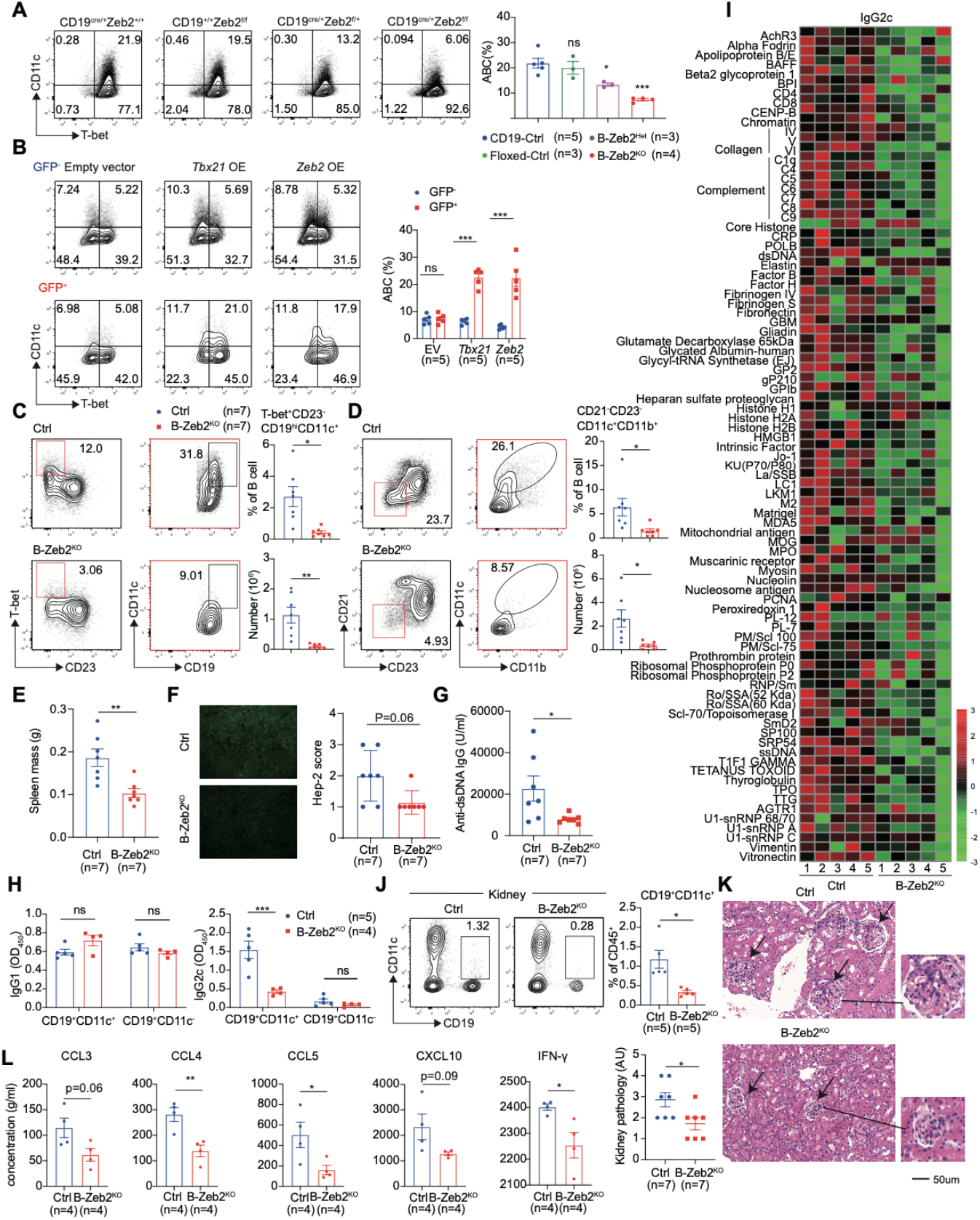
Zeb2 deficiency impairs ABC formation and alleviates lupus pathogenesis. (**A**) Splenic B cells were isolated from Zeb2^+/+^CD19^Cre/+^ (CD19-Ctrl), Zeb2^f/f^CD19^+/+^ (Floxed-Ctrl), Zeb2^f/+^CD19^cre/+^(B-Zeb2^Het^), Zeb2^f/f^ CD19^cre/+^(B-Zeb2^KO^) mice and cultured with ABC-skewing cocktail for 3 days. Representative plots and frequency of ABCs (CD19^+^CD11c^+^T-bet^+^). (**B**) Splenic B cells from C57BL/6J mouse were activated by R848, anti-CD40 and anti-IgM for one day, then retrovirally infected with overexpression plasmid pMYs-Zeb2 (Zeb2), pMYs-CTRL (EV), pMY-T-bet (T-bet) and further cultured with the ABC-skewing cocktail for further 3 days. Representative plots and frequency of ABCs (CD19^+^CD11c^+^T-bet^+^) in GFP^+^ and GFP^-^ B cells. (**C-D**) B-Zeb2^KO^ and CD19^Cre/+^ (Ctrl) mice were induced by imiquimod (IMQ) for 6 weeks. Flow cytometry plots and statistical analysis of splenic ABCs (CD19^+^CD23^-^CD11c^+^T-bet^+^, **C** and CD19^+^CD21^-^CD23^-^CD11c^+^CD11b^+^, **D**). (**E-G**) Spleen weight (**E**), ANA (**F**) and anti-dsDNA (**G**) in serum from mice described in (**C-D**). (**H**) Splenic CD19^+^CD21^-^CD11c^+^ and CD19^+^CD21^+^CD11c^-^ B cells sorted by flow cytometry from mice described in (**C-D**) were stimulated with anti-IgM, anti-CD40, R848, IL-21 for one day and then the supernatants were collected to detect IgG1 and IgG2c antibodies. (**I**) Autoantigen microarray was used to detect IgG2c autoantibodies in the serum of mice described in (**C-D**) by OmicsArray. (**J**) Representative plots and frequency of renal ABCs (CD19^+^CD11c^+^) from mice described in (**C-D**). (**K**) H&E staining (right) analysis of kidneys from mice described in (**C-D**), accessing kidney pathology (left) according to proliferation of glomerular mesangial cells and mononuclear cell infiltration. (**L**) Splenic CD19^+^CD21^-^ CD11c^+^ and CD19^+^CD21^+^CD11c^-^ B cells described in (**H**) were stimulated with anti-IgM, anti-CD40, R848, IL-21, and IFN-γ for one day and then the supernatants were collected for detection. n represents distinct samples (biological repeats). Data are representative of 2-3 independent experiments. Data are mean ± SEM values. *p<0.05, **P<0.01, ***P<0.001, ns, not significant, unpaired t-test (**B, E, H, K, L**) with Welch’s correction (**C-D, J**), Mann-Whitney test (**F, G**) and ordinary one-way ANOVA with Dunnett’s multiple comparisons testing (**A**).

Numerous studies in mice have demonstrated the physiological and pathogenic importance of ABCs in the humoral response, particularly of the IgG2a/c isotype (*1*). Indeed, targeting ABCs by depletion or genetic defects in mice can attenuate lupus pathogenesis (*2, 3, 5*). We thus interrogated whether Zeb2 regulated ABC-mediated autoimmunity. For this, we used a well-established lupus model mediated by chronic TLR7 signaling, by applying a low dose of the TLR7 agonist, imiquimod (IMQ) (*13, 39*). We confirmed ABCs (regardless of the markers used for identification: CD19^hi^CD11c^+^T-bet^+^CD23^-^ or CD21^-^CD23^-^CD11c^+^CD11b^+^ or CD19^hi^CD11c^+^CD23^-^) were significantly reduced in B-Zeb2^KO^ mice after IMQ treatment (Fig. 3, C and D, and fig. S6D). Automated clustering using tSNE algorithm for dimensionality reduction further confirmed that the frequency of ABCs, as a unique and discrete population identified by multiple markers, was dramatically decreased in B-Zeb2^KO^ mice (fig. S6, E and F). ABCs predominantly represent antigen-experienced cells with the capacity to differentiate into plasma cells (*1*). Our investigation demonstrated that a significant proportion of CD19^hi^CD11c^+^IgD^-^ ABCs exhibited CD38^+^GL-7^-^ memory markers, while also displaying a unique hyperactivated state (CD95^+^CD80^+^) in comparison to other memory B cells (Fig. S7A). Zeb2 deficiency selectively impacted the distribution and hyperactivation of CD11c^+^ memory-like B cells, without affecting the frequency of CD11c^-^ memory B cell subset (Fig. S7B).

Our study noted that a subset of ABCs presented a phenotype consistent with precursors of germinal center B cells (pre-GCB), indicating that during sustained chronic inflammation in the IMQ model, ABCs may be predisposed to enter the germinal center (GC) for the purpose of replenishing the GC pool and participating in ensuing chronic GC responses (fig. S7C). In B-Zeb2^KO^ mice, during chronic induction, the frequencies and absolute numbers of germinal center B cells and plasma cells were also diminished, albeit to a lesser extent than in ABCs, likely due to the reduction in the ABC-derived component (fig. S7, D to F). To further elucidate the effector B cell response, we utilized bm12-induced and LCMV infection models in B-Zeb2^KO^ mice. ABC populations were consistently decreased in both models, yet in contrast to the chronic IMQ model, primary GC B cells exhibited an augmentation after LCMV infection or bm12 induction, resulting from Zeb2 deficiency (Fig. S8, A to E). Additionally, there was a subtle enhancement of low-affinity antibody titers to an ABC-irrelevant antigen after hapten-protein conjugate immunization (NP-CGG plus alum and LPS) in the presence of comparable GC responses, suggesting that ABCs are distinct from the typical extrafollicular plasmablasts seen in conventional responses to protein antigens (fig. S8, F and G). Together, these experiments suggest that Zeb2 selectively drives ABCs and their offspring, which although typically extrafollicular, may participate in the GC response in the context of chronic inflammation.

In the IMQ model, *Zeb2* deficiency in B cells ameliorated splenomegaly significantly (Fig. 3E) and decreased serum autoantibodies against ANA and dsDNA (Fig. 3, F and G). To exclude the influence of antibody production arising from other trajectories, we sorted ABCs and non-ABCs from IMQ-treated mice according to the expression of surface markers CD11c and CD21 and then cultured them for 24 hours. Analysis of supernatants revealed that ABCs from Ctrl mice secreted the highest amount of IgG2c isotype antibodies upon re-stimulation (Fig. 3H). These data confirmed the ABC subset contained the majority of IgG2c isotype-switched B cells with the potential to produce IgG2c antibodies. However, CD11c^+^CD21^-^ B cells sorted from B-Zeb2^KO^ mice were unable to produce comparable IgG2c antibodies even though they met the gating criteria (CD19^+^CD11c^+^CD21^-^) for ABC-like cells by flow cytometry (Fig. 3H). The secretion of IgG1 antibody was not changed by *Zeb2* deficiency (Fig. 3H). A broader screen of autoantibodies using autoantigen microarrays revealed B-Zeb2^KO^ mice produced much lower IgG2c isotype autoantibodies after IMQ-induction (Fig. 3I). Taken together, these experiments show that *Zeb2* deficiency in B cells prevents ABC-mediated autoimmunity.

In lupus nephritis, ABCs have been shown to be present in the kidney and their abundance closely correlates with tissue damage (*7, 40*). ABCs are known to possess an increased capacity to produce proinflammatory cytokines and chemokines such as IL-6, IFNγ, CXCL10, and CCL5 (*5*), which might contribute directly to *in situ* damage and also recruit inflammatory cells to the affected organs. We thus checked whether *Zeb2* deficiency could dampen end-organ damage. In the IMQ-induced lupus model, we found kidney-infiltrating ABCs were dramatically decreased (Fig. 3J and fig. S7G) and tissue damage was alleviated (Fig. 3K) in mice lacking *Zeb2* in B cells. Ex vivo culture of the residual CD11c^+^CD21^-^ B cells from B-Zeb2^KO^ mice demonstrated that they could not produce comparable quantities of ABC-specific chemokines like CCL5 and CXCL10 (Fig. 3L), reflecting Zeb2 is required for the proinflammatory properties of ABCs.

These results confirm that Zeb2 is required for mouse ABC formation both in vitro and in vivo and, most importantly, that the pathogenic function of ABCs in lupus depends on *Zeb2* expression in B cells. Thus, Zeb2 specifically orchestrates ABC-related B cell response.

### Zeb2 controls the lineage specification and cellular identity of ABCs

The mechanisms involved in *Zeb2*-mediated gene regulation are broad and cell-context specific (*30, 31, 33, 36*). To investigate the consequences of *Zeb2* regulation of gene transcription we performed RNA sequencing on sorted B cells with reduced *Zeb2* expression (sg-*Zeb2* (CD19^+^GFP^+^BFP^+^)) 4 days after in vitro activation, retroviral infection, and differentiation. The ABC signature genes were compared between in vitro-derived ABCs from *Zeb2* deficient B cells and multiple transcriptomes of ABC-like cells from ageing, infection and autoimmune settings by public RNA-seq datasets GSE92387 (*6*), GSE110999 (*4*), GSE81650 (*41*), GSE81189 (*42*), GSE99480 (*5*) and our RNA-seq datasets of bm12 induced T-bet reporter mice (*43*) and in vitro-derived ABCs. Shared transcriptional profiles of ABC signature genes including *Itgax, Itgam, Itgb2, Nkg7, Tbx21, Zbtb32* and *Fcer2a/Fcer2* (encoding CD23a, CD23) in most ABC-like cells, were reversely expressed after *Zeb2* deficiency (Fig. 4, A and B). Gene set enrichment analysis (GSEA) identified that *Zeb2*-deficient B cells lacked expression of the “ABC upregulated” gene set while it was enriched with genes that characterize the “ABC downregulated” gene set from the public dataset GSE99480 (Fig. 4, C and D).

**Figure 4.**
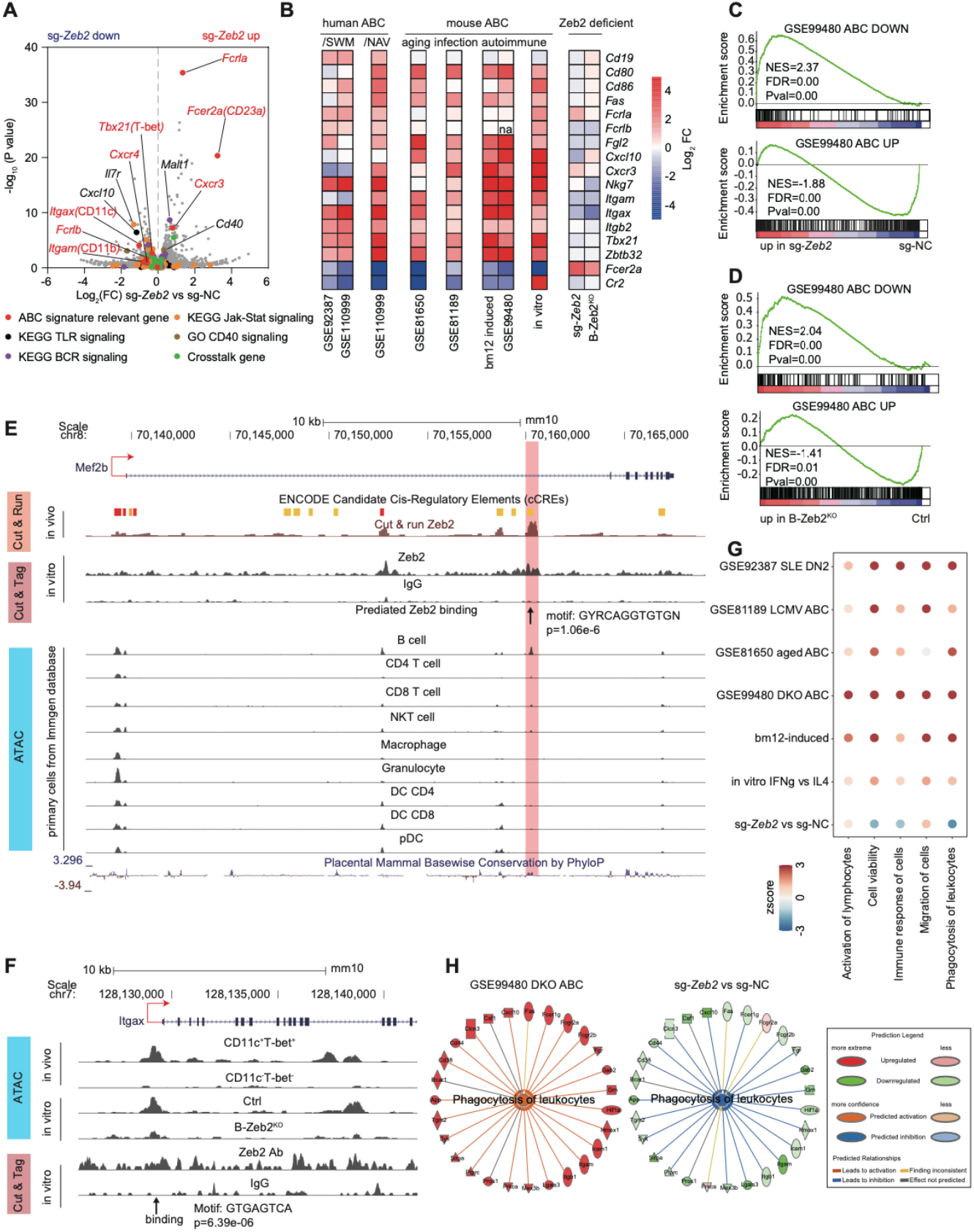
Zeb2 regulates specification and cellular identity of ABCs. (**A**) Volcano graph showing the transcriptional profiles in mouse B cells edited by sg-NC and sg-*Zeb2* under in vitro ABC induction for 3 days. ABC signature relevant genes, genes of BCR, TLR, Jak-Stat signaling, CD40 signaling and crosstalk genes were labeled with colored dots. (**B**) Heatmap showing expression of representative ABC signature genes in publicly available RNA-seq datasets: GSE92387, GSE110999, GSE81650, GSE81189, GSE99480 and our own RNA-seq datasets: bm12-induced (Splenic CD11c^+^T-bet^+^CD21^-^ and CD11c^-^T-bet^-^CD21^+^ B cells were sorted by flow cytometry from T-bet-tdTOMATO reporter mice immunized with bm12 T cells), in vitro (mouse B cells were cultured with ABC-skewing cocktail for 3 days), sg-*Zeb2* (described in (**A**)), B-Zeb2^KO^ (B cells from B-Zeb2^KO^ and Ctrl mice were cultured with ABC-skewing cocktail for 3 days). (**C-D**) GSEA showing the enrichment of the ABC-down geneset and ABC-up geneset from DKO mice (GSE99480) in sg-*Zeb2* versus sg-NC ABCs and B-Zeb2^KO^ versus Ctrl ABCs. (**E**) CUT & RUN, CUT & Tag, and ATAC-seq tracks display Zeb2 binding around the Mef2b locus in indicated immune cells, visualized with the UCSC genome browser. The chromatin accessibility of Mef2b in mouse primary immune cell subsets from Immgene database. (**F**) Chromatin accessibility by ATAC-seq and Zeb2-binding by CUT & Tag in the promoter site of mouse Itgax. ATAC-seq was performed in ex-vivo sorted ABC (CD11c^+^T-bet^+^) and non-ABC (CD11c^-^T-bet^-^) from bm12-induced mice and in-vitro derived Zeb2-deficient and ctrl B cell cultured for 3 day under ABC skewing condition. (**G**) Dot plot showing the activation Z-score for IPA-predicated biological function of ABCs including activation, cell viability, immune response, migration and phagocytosis from RNA-seq datasets described as in (**B**). (**H**) Network diagram representing phagocytosis pathway in sg-*Zeb2* versus sg-NC ABCs by IPA. The color of each node indicates change in the expression: red (upregulated) and green (downregulated). The edges indicate the predicted relationship between nodes and biological function: orange representing activation, blue representing inhibition, gray representing effect not predicted.

To identify the direct targets of Zeb2 regulation in ABCs, we performed a series of experiments. First, we examined the chromatin accessibility of Zeb2-driving ABCs to identify active regulatory regions and performed two transcription factor binding site characterization approaches: cleavage under targets and tagmentation (CUT & Tag) or release using nuclease (CUT & RUN), followed by sequencing the enriched DNA fragments to identify the binding sites of Zeb2 in ABCs (*44, 45*). We then combined the information from regulomes and found 600,002, 8,632, and 10,417 peaks from ATAC-seq, CUT & RUN, and CUT & Tag, respectively (fig. S9A). After overlapping, we identified 6,733 accessible sites with Zeb2 binding that could be annotated to 4,338 genes. Among the genes differentially expressed by Zeb2 deficiency, we found 33 candidate target genes of Zeb2, with 22 genes repressed and 11 genes activated by Zeb2 (fig. S9A).

One of the candidate target genes identified was Mef2b, which is a critical transcriptional factor for germinal center development (*46*) and was found to be repressed by Zeb2. We identified Zeb2 binding to a region about 20kb downstream of Mef2b exon I TSS from the enriched peaks of CUT & Tag and CUT & RUN (Fig. 4E). Importantly, this region had a site with Zeb2 binding consensus sequence that was conserved between human and mouse (fig. S9B). The conserved region displayed B cell-specific accessibility and had prominent enhancer-associated features, enriched with H3K4me1, H3K4me3, and H3K27ac modifications both in human and mouse (fig. S9, C and D). We validated Zeb2 suppression of Mef2b expression by qPCR and confirmed the opposing expression patterns of Zeb2 and Mef2b in ABCs and GCBs from public dataset GSE150124 (*47*) (fig. S9, E and F). Mef2b can directly regulate S1pr2 expression to fine-tune germinal center confinement (*46*), and in line with this, we observed upregulated expression of S1pr2 in Zeb2-deficient B cells (fig. S9E). These results suggest that Zeb2 likely promotes ABC differentiation by directly suppressing Mef2b to constrain GCB cells, consistent with our observations of the primary germinal center response in the bm12-induced and LCMV infection models (fig. S8, C and E).

Additionally, using CUT & RUN and CUT & Tag assays, we detected Zeb2 signals to be enriched near the transcriptional start sites (TSS), especially within the promoter regions (fig. S10, A and B). Zeb2-specific peaks from CUT & Tag assay were matched to motifs of *Gata3, Fosl2*, and *Zeb2* (fig. S10C), which is consistent with Zeb2 Chip-seq data in cell lines from the Encode project (fig. S10D). Further screening for functional targets, we identified a Zeb2-specific peak residing in the promoter of Itgax, containing a Zeb2-binding sequence (Fig. 4F). We demonstrated that Zeb2 deficiency altered the chromatin accessibility of the ABC-specific opening in the *Itgax* promoter, further confirming that Zeb2 modifies the transcriptional activity of ABC signature.

CD11c, an important alpha-subunit member of beta2 integrins, can pair with the beta-subunit CD18 (encoded by Itgb2) to form heterodimeric cell surface receptors for immune cell adhesion and recruitment to tissues (*48*), which is a unique property of ABCs (*4, 7, 40*). By analyzing biopsy samples of patients with lupus nephritis (LN) and para-tumor controls, we found that CD11c^+^ B cells remarkably infiltrated the affected tissues of LN patients with a substantial frequency of 50%. Furthermore, IgD^-^CD27^-^CD11c^+^ ABCs comprised about 20% of total B cells (fig. S11, A and B). To validate whether Zeb2-CD11c regulatory axis promotes the migration of ABCs, we performed transwell and trans-endothelial assays to assess B cell migration. We demonstrated that ABCs exhibit an enhanced migratory capacity, which can be modulated by CD11c blocking and Zeb2 deficiency (fig. S11, C to G).

Taken together, our results suggest that Zeb2 plays a crucial role in orchestrating ABC specification by directly suppressing other effector B cell subset and promoting the ABC signature. The Zeb2-driven regulation of *Mef2b* and *Itgax* provides insights into the molecular mechanisms underlying ABC differentiation, while the Zeb2-CD11c regulatory axis offers new avenues for understanding ABC migration and tissue infiltration.

### Zeb2 regulates the distinct biological features of ABC

Delineating and quantifying cellular transcriptomes provide comprehensive insights to understand cellular identity and function, as molecular changes are closely related to the alteration of biological processes and functions. We then applied a knowledge-based Ingenuity Pathway Analysis (IPA) to gain insights into ABC biology from the public dataset GSE99480. Among the top 35 significantly increased predicted biological functions (table S6), ABCs appeared to share 5 pathways: enhanced capacity for cell viability, migration, activation, immune response, and phagocytosis/engulfment (fig. S12A). Selected transcripts with the most relationships were linked to biological function and represented as a network (fig. S12A). We also analyzed the transcriptomes of ABCs from human individuals, the bm12-lupus model, in-vitro induction, or other settings (infection and ageing) and found those same 5 biological functions commonly increased in ABCs (Fig. 4G). Pathway analysis of these transcriptomes revealed that the enriched pathways were linked to the 5 functional features of ABCs (fig. S12B). ABCs’ hyperactivation state, long-term survival capacity, and unique migration/distribution pattern have also been observed in published studies (*4, 6, 49*). Interestingly, ABCs exhibited a unique phagocytic feature as identified by enriched phagosome formation and Fc receptor pathways (fig. S12B) which are normally considered functions of myeloid cells. The ability of ABCs to both conduct typical B cell functions and co-opt myeloid markers like the integrins CD11c and CD11b, and FcγRs, as well as some effector molecules, like NKG7, Granzyme A, and Perforin-1 has been previously recognized (*2, 4*). Comparative analysis revealed that *Zeb2* editing in B cells dampened cell viability, immune response, and phagocytosis/engulfment (Fig. 4G). Transcripts linked to phagocytosis were confirmed to display opposite expression patterns in *Zeb2* edited B cells (Fig. 4H).

To substantiate the notion that ABCs possess a unique phagocytic capacity, we incubated splenic B cells with apoptotic thymocytes labeled with pHrodo, a pH-sensitive fluorophore that facilitates the monitoring of apoptotic cell internalization (fig. S12C). Our findings indicated that CD19^hi^CD11c^+^ B cells exhibited a markedly enhanced uptake of apoptotic cells, as evidenced by both a higher frequency of pHrodo^+^ fraction and increased signal intensity (fig. S12D). Fluorescence microscopy provided direct visualization of ABCs engulfing apoptotic cells (fig. S12E). Notably, we observed that Zeb2 deficiency attenuated this distinct phagocytic capacity when in vitro-derived B cells under ABC-skewing conditions were incubated with pHrodo-labeled apoptotic cells (fig. S12F).

Collectively, these results demonstrate that, in comparison to other B cells, ABCs exhibit distinct biological functions, many of them regulated by Zeb2.

### Zeb2-Jak-Stat axis governs ABC differentiation

To more precisely identify the signaling pathways by which Zeb2 drives ABC formation, we performed Upstream Regulator Analysis (URA) in IPA on both public and our own datasets described above. As expected, BCR, CD40, and TLR and their key components like NF-Kb, and AKT were predicted to be activated in ABCs (Fig. 5A). The regulatory effects of several cytokines including IFNγ, IL-10, IL-21 and their downstream Jak-STAT signals were predicted to be altered in ABCs with trends of activation of *STAT1, STAT3*, and *STAT4* and inhibition of *STAT6* (Fig. 5A and B). All these enriched regulatory effects were consistent with the findings from previous studies of ABC biology (*1*). Zeb2 deficiency attenuated the effects of known ABC regulators including CD40, TLRs, and STATs (Fig. 5, A and B). Representative target genes of *STAT1, STAT3, STAT4*, and *STAT6* known to be critical for ABC differentiation (*1*) were listed in ABCs from the public gene set GSE99480 (Fig. 5B). By contrast, *Zeb2* editing in B cells led to opposite expression patterns of these target genes and altered regulatory effects of STAT1/3/4/6 (Fig. 5B). KEGG analysis confirmed that the top pathway of downregulated genes in *Zeb2* edited B cells were related to Jak-Stat signaling (fig. S13A) and GSEA also revealed altered Jak-Stat signaling in these B cells (Fig. 5C). Signaling landscape changes downstream of cytokine receptor signaling via Jak-Stat were annotated and genes involved in Jak-Stat signaling with significant changes were listed in a heatmap (Fig. 5D). We also confirmed the genes altered by Zeb2 deficiency largely overlapped with the transcriptional program affected by inhibition of Jak-Stat signal (fig S13, B and C). Together, these results suggest that Zeb2 regulates Jak-Stat signaling to drive ABC differentiation.

**Figure 5.**
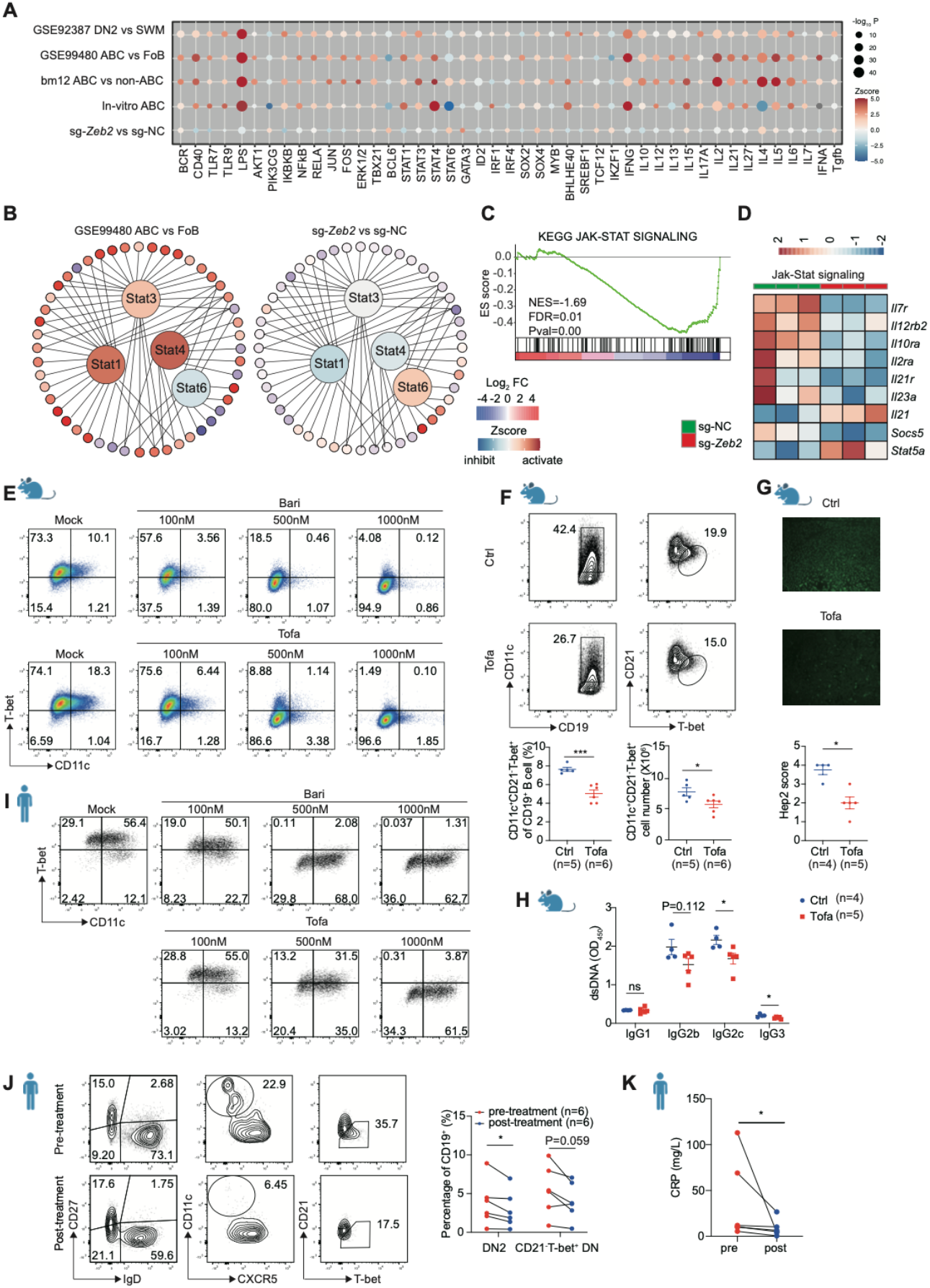
Zeb2-Jak-Stat axis controls ABC differentiation. (**A**) Activation *Z*-score heatmap for IPA-predicted upstream regulators in indicated datasets mainly described in **Fig 4B**. P-values was log-normalized and presented by the size of the plot. The color of the plot indicates the activated (orange) or inhibited (blue) regulation of the predicted regulators. (**B**) Upstream regulator analysis showing Stat1, Stat3, Stat4 activated and Stat6 inhibited in ABCs (DKO, GSE99480), while the opposite effects were observed in sg-*Zeb2* vs sg-NC dataset. The color of the surrounding circles indicates the change in the expression: red (upregulated) and blue (downregulated). The color of the center circles indicates predicted regulators: orange (activated) and blue (inhibited). (**C**) GSEA showing impaired Jak-Stat pathway in sg-*Zeb2*^+^ B cells compared with sg-NC. (**D**) Heatmap showing expression of selected Jak-Stat signaling genes in sg-NC^+^ and sg-Zeb2^+^ B cells. (**E**) Flow cytometry plots of in vitro induced mouse ABCs (CD11c^+^T-bet^+^) with different concentrations of baricitinib (Bari) or tofacitinib (Tofa). (**F-H**) C57 mice were immunized with bm12 T cells for 2 weeks and treated with tofacitinib (Tofa) or dissolved solution (Ctrl) by oral gavage once a day to analyze splenic ABCs (CD19^+^CD11C^+^T-bet^+^CD21^-^, **F**), ANA (**G**) and anti-dsDNA Ig (**H**) in serum. (**I**) Flow cytometry plots of in vitro induced human ABCs (CD11c^+^T-bet^+^) with different concentration of baricitinib or tofacitinib. (**J**) Flow cytometry plots of human ABCs (DN2, CD19^+^CD27^-^IgD^-^ CD11c^+^CXCR5^-^) or (CD21^-^T-bet^+^ DN, CD19^+^CD27^-^IgD^-^CD21^-^T-bet^+^) in PBMCs from patients with rheumatoid arthritis before and after Jak-Stat inhibitor tofacitinib treatment for 4 weeks. (**K**) CRP level in RA patients described in (**J**). n represents distinct samples (biological repeats). Data are representative of 2-3 independent experiments. Data are mean ± SEM values. *p<0.05, **P<0.01, ***P<0.001, ns, not significant, unpaired t-test (**F, H**), Mann-Whitney test (**G**), paired t-test (**J**) and paired Wilcoxon test (**K**).

Jak-Stat inhibitors have proven to be a safe and efficient strategy to dampen intracellular cytokine signaling in various cell types (*50, 51*). Baricitinib, tofacitinib, upadacitinib, and filgotinib have been approved for oral therapy in rheumatoid arthritis (RA) and are also commonly prescribed off-label in other autoimmune diseases including SLE, inflammatory bowel disease, and psoriasis (*51*). We therefore investigated the effects of Jak-Stat inhibitors on ABC formation. Baricitinib or tofacitinib impaired mouse in vitro ABC differentiation in a dose-dependent manner (Fig. 5E, and fig. S13, D to G). Oral administration of tofacitinib effectively reduced the accumulation of ABCs (Fig. 5F) and splenomegaly (fig. S14A), and also decreased autoantibody titers (ANA and anti-dsDNA) in the bm12-induced lupus model (Fig. 5, G and H). Given the critical role of the Jak-Stat pathway in both adaptive and innate immune cells, we sought to validate whether tofacitinib therapy could reduce inflammatory cytokine expression, including IL-6 and GM-CSF. Our results confirmed that tofacitinib treatment effectively decreased the expression of these cytokines but did not inhibit the generation of IFNγ and IL-21 which were required for ABC differentiation, nor the distribution of IFNγ-producing Th1 cells (fig. S14, B and C). These findings suggest that the reduced accumulation of ABCs by tofacitinib is likely due to B-cell intrinsic inhibition of the cytokine response, rather than indirect effects via altered cytokine production although this cannot be definitively excluded. Additionally, we observed a reduction in germinal center B cells and plasma cells by tofacitinib which could also contribute the therapeutic efficacy (fig. S14, D and E). But considering IgG2c isotype of anti-dsDNA antibodies was the most abundant and significantly decreased after tofacitinib treatment (Fig. 5H), this improved serological phenotype was more likely associated with impaired ABC accumulation than other B cells.

Accordingly, human ABC differentiation in vitro was also blocked by baricitinib and tofacitinib (Fig 5I, and fig. S14F). Finally, we investigated the *in vivo* effects of tofacitinib on ABC differentiation in RA patients. The frequency of human ABCs (DN2) had decreased 4 weeks after treatment (Fig. 5J and fig. S14G). Tofacitinib also ameliorated systemic inflammation lowering C-reactive protein (CRP) (Fig. 5K). These results demonstrate that besides the efficacy from other immune cells, targeting the Jak-Stat pathway is also an efficient strategy to block ABCs in mice and humans and provides a rationale for the use of Jak-Stat inhibition therapy in ABC-mediated autoimmunity.

## Discussion

Elucidation of the TF determining ABC differentiation is essential to understand the ontogeny and role of this B cell subset and to develop ABC targeted therapies particularly in the context of systemic autoimmunity. Our results have identified Zeb2 as an essential TF that drives human and mouse ABC differentiation and ABC-mediated autoimmunity including autoantibody formation, production of proinflammatory cytokines and chemokines, migration to end-organ tissues, and tissue damage.

Zeb2 restricts GCB differentiation via direct Mef2b repression while promoting ABC identity by directly targeting signature genes including *Itgax*. Transcriptional regulatory networks have been established for early and late B cell lineage development (*52, 53*). It is likely that there is also a stepwise regulatory program for ABC specification and commitment. For instance, IRF5 directly promotes cytokine production and IgG2a/c isotype switching (*5*), whereas T-bet is also required for IgG2a/c isotype switching (*54*) as well as for terminal differentiation to plasma cells (*10*) and metabolic control of ABCs (*43*). Zeb2 is likely to act in concert with these and other TFs or epigenetic factors to orchestrate a regulatory complex in ABCs, similar to the regulatory programs of NK cells (*55*) and CD8 T cells (*33*) where Zeb2 is also involved.

Besides driving ABC differentiation, Zeb2 also determines important biological features of this subset. Here we show that ABCs can phagocytose apoptotic cells, which is a feature of innate immune cells. Internalization of apoptotic cells and other cellular debris may provide a source of nucleic acid ligands for TLR7 activation and self-antigens for presentation to T cells, as well as explain the hyperactivated status of this B cell subset.

Mice and humans have ABC subsets that share a similar phenotype and exert pathogenic roles in autoimmunity (*1*). It is therefore not surprising that both require Zeb2 for ABC development. Based on our observation that Zeb2 regulates ABCs at least in part via the Jak-STAT pathway, we rationalized that therapeutic intervention with Jak inhibitors would be beneficial for ABC-mediated autoimmunity. Indeed, this turned out to be the case in both mice and humans. Although our study has focused on the impact of Zeb2-driven ABC formation in the context of autoimmunity, our results are likely to apply to other settings in which ABCs have been shown to be expanded such as microbial infections and ageing.

## Acknowledgments

We thank Z. Liu, Y. Ma, Q. Hu, Y. Hu, Y. Chen and S. Zhou for providing experimental help; J. Li, X. Xu for provding the LCMV Amstrong virus; X. Song for suggestion; J. Qin and J. Huang for sample collection and Y. Yu for reagents support.

## Funding

National Natural Science Foundation of China (31630021, 31930037, 82071843 and 81901637)

National Human Genetic Resources Sharing Service Platform (2005DKA21300)

Shanghai Municipal Key Medical Center Construction Project (2017ZZ01024-002)

Shenzhen Science and Technology Project (JCYJ20180504170414637 and JCYJ20180302145033769)

Shenzhen Futian Public Welfare Scientific Research Project (FTWS2018005)

Sanming Project of Medicine in Shenzhen (SZSM201602087).

## Author contributions

Conceptualization: NS, DD, CGV

Methodology: DD, SG

Investigation: DD, SG, XH, HD, YJ, XZ, CY, SH, JZ, GH, BQ, HZ, YQ, YH, JM, ZY, ZY, JQ, QJ, LW, QG, SC, CH, LCK, MTW

Visualization: DD, SG, YJ

Funding acquisition: NS, DD, ZY, SC

Supervision: NS, CGV

Writing-original draft: DD, GSS

Writing-review & editing: NS, CGV

## Data and materials availability

All data are available in the main text or the supplementary materials. The sc-RNAseq, RNAseq, ATAC-seq, Cut & Tag and Cut & Run sequencing data have been deposited in ArrayExpress database and the accession code will be provided prior to publication. Source data are available upon request. Plasmids are available upon establishment of an MTA with Shanghai Jiaotong University.

## Supplementary Materials

Materials and Methods

Figs. S1 to S14

Tables S1 to S9

Reference (*56-69*)

