## supplementary figures and tables for "The transcription factor Zeb2 drives formation of age-associated B cells"

### Materials and Methods

#### Mice, bm12 cGVHD, imiquimod (IMQ) lupus models, LCMV infection, 4-Hydroxy-3-nitrophenyl (NP) acetyl-hapten immunization model

The C57BL/6J (JAX000664), bm12 (JAX001162), Cd19-Cre (JAX006785), B6 Cd45.1 (JAX002014) and Cas9-EGFP (JAX026175) mice were obtained from Jackson Laboratory. Zeb2$^{flox}$ mice (T015122) and T-bet-tdtomato (T036727) (*43*) mice were obtained from Gempharmatech (Nanjing). All the mouse strains were on C57BL/6J background and maintained under specific pathogen free (SPF) conditions in Renji Hospital animal facilities. All procedures involving animals were approved by Institutional Animal Care and Use Committee at Renji Hospital of Shanghai Jiaotong University School of Medicine (SJTUSM). Mice were randomly divided into groups and injected by (H.X.) while sample acquiring was conducted by (G.S.), Data analysis was performed by (D.D.). No samples were excluded from analysis.

bm12-cGVHD lupus model was induced following JOVE protocol (*56*). Briefly, 8-12-week-old mice were intraperitoneally injected with 10 million CD4$^+$ T cells (Miltenyi Biotec, Cat#130-104-454) from bm12 mice purified by magnetic cell sorting and mice were sacrificed for analysis at indicated time points. For Jak-Stat inhibitor treatment experiment, mice were orally gavaged with 30 mg/kg tofacitinib (Selleckchem, Cat#S2789) for 2 weeks.

IMQ-induced lupus model was induced following previous protocol (*39*). Briefly, 8-12-week-old mice were treated topically with 1.25mg of 5% imiquimod cream (3M Health Care Limited) applied to the suitable ear skin three times a week and mice were sacrificed for analysis at indicated time points. Mouse kidneys were fixed in 4% neutral buffered formalin and embedded in paraffin. Sections were stained with H&E and were analyzed by light microscopy. The kidney pathologic score accounted for the morphological pattern (glomerular mesangial cells and inflammatory cellular infiltration) using Case Viewer software (3DHistech).

NP immunization model was induced by NP-CGG plus alum and LPS as adjuvant. Briefly, alum was dropwise added to NP-CGG solution (containing 1mg/ml NP-CGG and 10ug/ml LPS) at a 1:1 volumetric ratio while vortexing. 8-12-week-old mice were intraperitoneally administered 200ul of NP-CGG/Alum/LPS. After induction for 14 days mice were sacrificed for analysis.

LCMV infection was induced by intraperitoneally injection of $1.5 \times 10^5$ PFU of LCMV-Armstrong for 10 days and mice were sacrificed for analysis.

#### Patients subject

SLE and RA patients were recruited from Renji Hospital. SLE patients met American College of Rheumatology criteria for SLE. Demographic and clinical information is listed (table S1 for scRNAseq, table S2 for patients with lupus nephritis for collecting biopsy samples). RA patients met the 2010 American College of Rheumatology criteria for RA. Clinical and laboratory data of RA patients were measured at baseline and at week 4 after tofacitinib treatment. Demographic and clinical information is listed before and after treatment (table S3).

MWS patients were recruited from Children's Hospital, Capital Institute of Pediatrics. Patients were diagnosed based on a thorough clinical evaluation, identification of characteristic physical findings and facial appearance and confirmed by genetic testing for mutations in the *ZEB2* gene. Demographic and clinical information is listed (table S4, S5).

Written informed consent was obtained from all patients, healthy controls, or their parents. All procedures were fully consent under ethical and safe protocols approved by the Review Board at Renji Hospital of SJTUSM and Children's Hospital, Capital Institute of Pediatrics.

**Cell isolation, culture flow cytometry and sorting**
Single cell suspensions of splenocytes were subjected to red blood cell lysis and filtered through 70 um nylon mesh. Cells were stained for surface markers or intracellular proteins under instructions (*57*). For in vitro differentiation experiment, B cells transfected sgRNA plasmids (BFP) were stained with surface markers and then fixed by 2% fresh prepared formaldehyde in PBS for 5 minutes to allow efficient TF and cytokine staining and retain FP which were optimized from previous protocol (*58*). For intracellular T-bet staining, cells were fixed and permeabilized after surface staining using Foxp3 Transcription Factor Staining Buffer Set (ThermoFisher, Cat#00-5523-00). Samples were acquired using LSRFortessa (BD Biosciences) and analyzed using FlowJo software (Tree Star). Gating strategies for flow cytometry plots were displayed (fig. S2, A and B, fig. S3, A and B, fig. S5, A and C, fig. S6, D and F, fig. S7, D and G, fig. S14G).

To isolate kidney infiltrated cells, mice kidney tissues were minced and digested with type IV collagenase (250ng/ml, sigma, Cat # C5138) and DNase I (10U/ml, sigma, Cat # D5025) for 30min at 37°C in a shaking incubator. Cells were filtered through a fine 70 μm nylon cell strainer, suspended in 40% Percoll underlaid with 80% Percoll (GE Healthcare Life Sciences, Cat#17-0891-01), and lymphocytes were enriched in the middle layer using density-gradient centrifugation. Cells were harvested and washed for further analysis.

For sorting experiment, splenic B cells were enriched by magnetic microparticle purification using mouse B Cell Isolation Kit (Miltenyi Biotec, Cat#130-090-862) and then further sorted by gating $CD11c^+$ B cells ($CD19^+CD11c^+CD21^-$) and $CD11c^-$ B cells ($CD19^+CD11c^-CD21^+$) using FACSAria II (BD Biosciences) (fig S2B). Detailed information of antibodies were listed in table S7.

To ex vivo culture $CD11c^+$ B cells ($CD19^+CD11c^+CD21^-$) and $CD11c^-$ B cells ($CD19^+CD11c^-CD21^+$) from IMQ-induced lupus model, sorted cells were cultured with complete RPMI-1640 medium at a concentration of $1x10^6$ cells/ml for 24h and stimulated in 96-well plates with R848 (500ng/ml), anti-CD40 (1ug/ml), IL-21 (50ng/ml) and/or IFN-γ (10ng/ml). Then the culture supernatants were collected to detect antibodies, cytokines and chemokines.

**T-SNE visualization of flow cytometric data**
All samples were pre-gated on live single lymphocytes and each sample was randomly down sampled to 5000 events and merged into a single expression matrix. T-Stochastic Neighbor Embedding was applied to reduce the dimensionality of the data by using the T-SNE plugin available on the FlowJo Exchange. The composite sample then was hand-gated as indicated for all populations to aid in visual overlays with T-SNE maps. Intensities for markers of interest were overlaid on the T-SNE maps to show the expression of those markers on different cell clusters.

**In vitro induction of ABC**

Mouse splenic B cells were purified by negative selection and cultured in RPMI-1640 medium with 10% fetal bovine serum, HEPES, non-essential amino acids (NEAA), glutamine, sodium pyruvate, β-Mercaptoethanol and penicillin/streptomycin. ABCs were induced in the presence of R848 (500 ng/ml, Invivogen, Cat#tlrl-r848-5), anti-CD40 (1 ug/ml, Biolegend, Cat#102812), anti-IgM F(ab')$_2$ (1 ug/ml, Jackson Immunoresearch, Cat#115-006-020), IL-21 (50 ng/ml, Peprotech, Cat#210-21), IFN-γ (1, 5 or 10ng/ml, Biotech, Cat#485-MI-100) for 3 days.

Human PBMCs were isolated by density gradient centrifugation using Ficoll-Paque (GE Healthcare, Cat#17-5442-03). B cells were purified by negative selection (Miltenyi Biotec, Cat#130-091-151) from PBMCs and stimulated with R848 (1 ug/ml), CD40L (10 ug/ml, Biolegend, Cat#591708), BAFF (20 ng/ml, Peprotech, Cat#310-13-20), IL-2 (10 ng/ml, Peprotech, Cat#200-02), Goat Anti-Human IgA + IgG + IgM (H+L) (10 ug/ml, Jackson Immunoresearch, Cat#109-006-064), IL-21 (10 ng/ml, Biolegend, Cat#571204), IFN-γ (20 ng/ml, Biolegend, Cat#570206) for 3 days and stained with anti-human CD19, CD38, CD27, IgD, CD11c, CD21, T-bet to detect ABC formation. Addition of Tofacitinib and Baricitinib (Selleckchem, Cat#S2851) were indicated.

**In vitro apoptotic cell phagocytosis assay**

To induce apoptotic cells, thymocytes were collected from C57BL/6 mice and cultured in RPMI-1640 medium containing 10% FBS, 1% penicillin/streptomycin, and Camptothecin (500nM) for 24 h. The efficiency of apoptosis induction was verified by an apoptosis detection kit (containing Annexin-V/Propidium Iodide) according to the manufacturer's instructions with above 80% apoptotic cells (Biolegend, Cat#640932).

Then the apoptotic thymocytes were labeled with 60 ng/ml pHrodo (Invitrogen, Cat#P35357) for 2 h at 37 °C. Labeled apoptotic thymocytes and target cells (splenocytes from lupus mice or in vitro derived B cells) were mixed at a 10:1 (apoptotic thymocytes: target cells) and incubated in the complete RPMI-1640 medium with R848 (1ug/ml, Invivogen, Cat#tlrl-r848-5) for 120 min. Phagocytosis of the labeled apoptotic thymocytes was then evaluated by flow cytometry.

**Time-lapse microscopy assay for phagocytosis**

To conduct the real-time cellular phagocytosis analysis, the IncuCyte® pHrodo® Red Cell Labeling Kit (Sartorius, Cat#4649) was used for cell labeling and Incucyte® Live-Cell Analysis System was used for images collection. Apoptotic thymocytes were washed by IncuCyte® pHrodo® Wash Buffer at first step and then resuspended in IncuCyte pHrodo Labeling Buffer. The cells were labeled with IncuCyte pHrodo Red Cell Labeling Dye (100 ng/ml) at a cell concentration of $1x10^6$/ml for 60 min at 37 °C. After incubation, pHrodo-labeled cells were washed and resuspended in complete medium. $5X10^6$ labeled apoptotic thymocytes were then mixed with $5x10^5$ ABCs sorted from lupus mice at a 10:1 ratio (apoptotic thymocytes: ABC) and incubated in complete RPMI-1640 medium with R848 (1 ug/ml, Invivogen, Cat#tlrl-r848-5) in 24 well Plate. The plate was transferred to the Incucyte® Live-Cell Analysis System and each well was imaged for 4 pictures every 30 minutes.

**In vitro Transwell migration assay and Transendothelial migration assay**

To detect transwell migration, B cells were isolated from spleen of B-Zeb2$^{ko}$ mice and control mice and cultured under ABC skewing condition for three days. In vitro–induced B cells were purified using Dead Cell Removal Microbeads (Miltenyi Biotec, Cat#130-090-101) and then resuspended with migration medium (RPMI 1640 supplemented with 0.5% fatty acid–free BSA, 1% penicillin/streptomycin, and 1% Hepes).

Transwell insert (5μm, Corning; Cat #CLS3421) was placed on top of each well. 100 μl of cells (6 X$10^5$ in vitro derived B cells and 4X$10^5$ primary CD45.1$^+$ B cell) were added into the upper chamber and allowed to migrate for 6h, after which migrated cells in the lower chamber were collected and analyzed by flow cytometry. To explore the role of CD11c during ABCs migration, B cells isolated from lupus mice were cocultured with the purified anti-mouse CD11c Antibody (25 ug/ml, Biolegend, Cat# 117302) for 1h and then prepared for transwell migration assay as described above.
For trans-endothelial migration assay, transwell inserts were coated with Matrigel (Corning, Cat#356324) for 30 minutes at 37°C. Then 5X$10^4$ murine endothelial cell line (bEnd.3) were prepared and seeded on the upper surface of transwell insert that had been previously coated with Martrigel. bEnd.3 cells were cultured for 24 hours to form a compact monolayer, and then B cells from lupus mice were added to the upper chamber of each Transwell. The lower chamber was added with CXCL12 (100 ng/ml, Peprotech; Cat # 25020B). After a 6h incubation at 37°C, cells in the lower chamber were collected and analyzed by flow cytometry.

**Vector construction, sgRNA cloning, retrovirus packaging and infection**
For the transcription factor screening, sgRNA designed by online design tool (http://chopchop.cbu.uib.no/) and two DNA oligo annealing by T4 Polynucleotide Kinase (NEB, Cat#M0201L) and cloned into a Bbs1-digested U6-sgRNA-PGK-Puro-BFP retrovirus vector by T4 DNA ligation. The day before transfection, 80 million Plate-E cells were plated in 10 cm dish in DMEM with 10% FBS. Cells were transfected with 15 ug sgRNA plasmid using 30 μL of Lipofectamine 2000 (Invitrogen, Cat#11668019). Plat-E cells were further cultured for 48 hours. The retrovirus supernatants were centrifuged and debris were removed. Retrovirus were spin transduced to activated B cells in the presence of polybrene (10 ug/ml, Sigma, Cat#H9268) with 1260 RCF for 90 minutes at temperature of 32℃. The medium was removed six hours later and replaced with fresh medium for ABC induction. At least 4 single guide RNA (sgRNA) plasmids (co expressing blue fluorescent protein, BFP as reporter) were constructed per gene target and validated editing efficiency in mouse primary B cells using the online ICE tool (Inference of CRISPR Edits; Synthego) (*59*). We select the most efficient sgRNA with an editing rate over 50% for further functional screening. The selected sgRNA sequences were listed in **table S8**.

To overexpress Zeb2 and T-bet, the overexpression plasmids were constructed by Tsingke Biotechnology Co.Ltd. (Shanghai, China). Simply, cDNA sequences of mouse Zeb2 and Tbx21 gene cloned by PCR were inserted into pMYs-IRES-GFP vectors and then were verified by sequencing. Retrovirus described above was transduced to activated B cells in ABC skewing cocktails for 3 days to further analyze ABC frequency by flow cytometry.

**Human B cell editing**
Purified B cells from human PBMC were stimulated with R848 (1ug/ml), IL-2 (10ng/ml), BAFF (20ng/ml), Goat Anti-Human IgA/IgG /IgM (H+L) (10ug/ml) for one day. The next day, synthetic

sgRNA oligos (GenScripts) were co-incubated with Cas9 protein (GenScripts, Cat#Z03389) at room temperature for 20 minutes. 0.1 million cells were washed and electroporated using the Neon transfection system (Thermofisher, Cat#MPK5000S) following manufacturer's instruction. For candidate TFs, we selected sgRNAs with editing efficiency over 50%. The selected sgRNA sequences were listed in **table S8**. After electronical transfection, human B cells were induced to ABC differentiation described above.

**Antibody titers, autoantigen array and multiplex Immunoassay assay**
Serum samples were collected at indicated time points to detect anti-dsDNA by ELISA (Alpha Diagnostic International, Cat#5110) and Hep-2 cells slides (Inova diagnostics Cat#708750) were used to detect anti-nuclear antibodies. FITC-anti-mouse IgG were used as detection antibody for ANA. Antibody from culture supernatant was diluted and captured by goat anti-mouse Ig (Southern Biotechnology, Cat#5300-05B) and detected by HRP- labeled goat anti-mouse IgG1, IgG2c, IgG3 or IgM specific Abs and TMB substrate (Biotech, Cat#DY999) followed by sulfuric acid stop. Absorbance values at 370/450 nm (OD) were read and absolute titers were calculated from standard curves, which were generated from purified mouse Ig isotype (Southern Biotechnology, Cat#5300-01B).

To profile IgG2c isotype autoantibodies in the serum of B-Zeb2$^{KO}$ and Ctrl mice, reactivities against a panel of 128 autoantigen specificities were measured using an Autoantigen Microarray platform developed by the Genomics and Microarray Core Facility, UT Southwestern Medical Center.

To detect the cytokine and chemokine production, culture supernatant was collected and detected by Bio-Plex Pro Mouse Chemokine Panel 31-Plex (Bio-Rad, cat#12009159) according to manufacturer instructions.

**RNA extraction and quantitative RT-PCR analysis**
Total RNA was extracted from B cells with TRIzol reagent (ThermoFisher, Cat#15596026) according to the manufacturer's instructions. RNA quantity and quality were assessed using the Nanodrop 2000. cDNA was synthesized using PrimeScript™ RT Reagent Kit (TaKaRa, Cat#RR037A), Real-time PCR reactions were performed using TB Green Premix Ex Taq reagent (TaKaRa, Cat#RR420A) on QuantStudio 7 Flex (Applied Biosystems). For quantification of gene expression, each sample was normalized to expression of an endogenous control gene Rpl13a. All primers used were listed in **table S9**.

**Single-cell RNA library preparation and sequencing**
After FACS sorting of ZA$^{-}$CD19$^{+}$ B cells from PBMC of patient with new-onset SLE, 12000 cells were counted and viability was confirmed to be >90%. Single-cell suspensions were loaded on a 10X Genomics Chromium single-cell 5′ v2 chip to prepare the libraries according to the manufacturer's protocol. The library quality was checked by Agilent 2100 Bioanalyzer and Qubit fluorometric quantitation. Samples were sequenced on an Illumina NovaSeq 6000 with a sequencing depth of at least 50k reads per cell.

**Processing of single-cell RNA-seq data**

The raw FASTQ data were processed with Cell Ranger (v3.1.1) (*60*) by using GRCm38 annotation (v1.2.0). Output from Cell Ranger was loaded into R and used Seurat (v3.1.1) (*61*) package for further analysis. Total 5721 cells with fewer than 500 or more than 5000 expressed genes, a high percentage of mitochondrial genes (>10%) were removed. Next, the UMI counts in each cell were normalized and scaled. The top 1500 highly variable genes were used for dimensionality reduction using Principal Components Analysis. Upon 'Elbow plot', we selected the first 15 PC for clustering and UMAP visualization with default parameters. For clustering, we first selected resolution parameter = 0.8 which produced 9 clusters. To obtain cluster-specific gene signatures, we identified differential expression analysis of each cluster against the others using wilcox with parameter (logfc. threshold = 0.25, min.pct=0.1). Clusters were manually merged with the nearest cluster based on the phylogenetic tree from Seurat's Build Cluster Tree and similar marker genes. Finally, we got 7 clusters for downstream analysis.

**RNAseq**

Cas9-EGFP mouse B cells were retrovirally transduced with sg-NC, sg-Zeb2, sg-T-bet plasmids after activation for 1 day and further cultured under in vitro ABC induction condition for 72 hours. Cells were stained with anti-mouse CD19 and $CD19^+EGFP^+BV421^+$ cells were sorted. Splenic B cells were in vitro induced for ABC formation as describe above with or without tofacitinib (1000nM) treatment. Cells were washed twice and resuspended in TRIzol reagent. Total RNA was extracted and used as input for Illumina TruSeq RNAseq library kit. After quality assessment, RNAseq libraries were pooled. 150 bp paired-end sequencing was performed using Illumina Novaseq6000.

Clean reads were aligned to mm10 reference genome with Hisat2 (v2.1.0) (*62*) using default parameters. Gene expression levels were counted with HT-seq (v0.11.2) (*63*) (*64*) using default 'union' mode. Differential expressions of genes were conducted by DESeq2 (v1.24.0) package (*65*) and simple plots were produced in R (v3.3.3). GSEA was performed using the online software (Broad Institute) (http://software.broadinstitute.org/gsea/index.jsp).

**ATAC-seq and data analysis**

ATAC-seq was performed with TruePrepTM DNA Library Prep Kit V2 for Illumina®（Vazyme, Cat#TD501). $2X10^4$ $CD11c^+T\text{-}bet^+$ and $CD11c^-T\text{-}bet^-$ B cells were sorted by flow cytometry from T-bet reporter mice. B-$Zeb2^{KO}$ and Ctrl B cells were cultured under the ABC-skewing system for 3 days as described above. Collected cells were resuspended in lysis buffer (10mM Tris-Cl, 10mM NaCl, 3mM $MgCl_2$, 0.1%Igepal CA-630) and incubated on ice for 10min following centrifuge. The cell pellet was resuspended in transposase reaction mix (10ul 5XTTBL, 5ul TTE Mix V50, 35ul nuclease-free water) and incubated at 37 °C for 30 min. Fragmented DNA was purified using VAHTS DNA Clean Beads (Vazyme, Cat#N411-01) and library was generated by PCR using TruePrep® Index Kit V3 for Illumina® (Vazyme, Cat#TD203) for 11 cycles. PCR cleanup of libraries was performed using VAHTS DNA Clean Beads at a 1:1.2 ratio. Libraries were then sequenced on Illumina NovaSeq 6000 with paired-end reads.
Sequencing data were analyzed as follows. Data were mapped to the mouse reference genome (GRCm38/mm10) using Bowtie2 with the default setting followed by removing PCR duplicates using Sambamba. Peaks were called using Genrich with -FDR 0.01. For track display, alignments were converted to bigwig file using bedtools.

**CUT&RUN and data analysis**

CUT&RUN libraries were generated following manufacturer's instructions (CST, Cat#86652). Briefly, $5X10^5$ cells $CD19^{hi}CD11c^+CD21^-$ cells were sorted from lupus mice and incubated with CoA-coated magnetic beads for each sample. Cells were permeabilized and then incubated with primary antibodies against Zeb2 and IgG for 2 hrs following with secondary antibody. The beads-cells mixture was resuspended in protein A MNase and incubated for 1h. Then, the liquid was transferred to a new tube for phenol-chloroform-isoamyl alcohol DNA extraction. Libraries were cleaned up using VAHTS DNA Clean Beads (Vazyme, Cat#N411-01).

The sequencing libraries were prepared with the NEBNext Ultra II DNA Library Prep Kit for Illumina (NEB, Cat#E7645S) according to manufacturer's protocol. Libraries with different indexes from NEBNext® Multiplex Oligos for Illumina® (NEB, #E7335) were pooled and sequenced with an Illumina Hiseq-PE150 as paired-end reads extending 150 bases.

Sequencing data were analyzed as follows. The adaptor sequences were discarded by Trimmomatic before alignment. To visualize CUT&RUN datasets with UCSC genome browser, the CUT&RUN reads were aligned and mapped to the mouse reference genome (GRCm38/mm10) by Bowtie2 software. Duplicated reads were removed using 'make tag directory' in Homer with the parameter -tbp 1. Peaks from individual conditions were identified by findPeaks in Homer and the resultant files were converted to BED files by pos2bed.pl in Homer for visualization in UCSC genome browser.

**CUT-Tag and data analysis**

CUT&Tag libraries were generated following instructions described previously (*66*) and manufacturer's protocol (Vazyme, Cat#TD901-01). Briefly, 0.5 million in vitro derived mouse ABCs were incubated with CoA-coated magnetic beads, and then primary antibodies (anti-Zeb2) were added to incubate for 1 hours at room temperature. After incubation, secondary antibodies (incubated at RT for 1 hour) and pG-Tn5 adapter complex (incubated at RT for 1 hour) were added. Then, tagmentation took place for 1 hour, and DNAs were extracted using phenol–chloroform–isoamyl alcohol. Libraries were prepared and cleaned up using Ampure XP beads (Beckman Coulter) and pooled together for paired-end sequencing.

Sequencing data were analyzed as follows. After trimming the adapters of reads and removing low-quality bases by TrimGalore software in paired-end mode, clean reads were mapped to mm10 reference genome with Bowtie2 (v2.3.5) (*67*) followed by removing PCR duplicates using Picard (v2.19.0).The genome browser tracks in bigwig format were produced from merged replicates using deepTools (v2.0) (*68*) and peaks were called using MACS2 (v2.1.2) (*69*) using parameters '-f BAMPE –SPMR -nomodel'. Enriched transcription factor binding motifs were searched by HOMER (v4.11). A set of sequences were searched for individual matches to each of the motif using FIMO (https://meme-suite.org/meme/tools/fimo). All analysis data were visualized using the UCSC genome browser.

**Peaks overlap and annotations**

Intersect in bedtools was used to screen for overlaps among peaks from ATAC-seq of $CD11c^+T\text{-}bet^+$ B cells, Zeb2 CUT&RUN in $CD19^{hi}CD11c^+CD21^-$ cells, and Zeb2 CUT&Tag in in vitro derived mouse ABCs as described above. Annotation of the overlapping peaks were performed with ChIPseeker.

**IPA analysis**

To define the distinct biological function and upstream regulators of ABCs, we integrated the public datasets and our own datasets to perform core analysis using Ingenuity Pathway Analysis (IPA) software. The cut-off for ABC specific genes from different settings was set as expression $\log_2$ fold change ≥1 or ≤−1 and statistically significant FDR value <0.05. The cut-off for sg-*Zeb2* vs sg-NC was statistically significant FDR value <0.05. The significance and regulatory effects of enriched pathways, upstream regulators and biological functions was qualified using overlap P value and activation Z score based on the observed pattern of up/downregulation of the target molecules compared with expected directions of changes documented in Ingenuity's curated database.

**Statistical analysis**

Prism software (GraphPad) was used for statistical analyses. Data were shown as mean ± SEM values. All data points represented the measurement of distinct samples. Statistical tests were selected based on the distribution and the variance characteristics of the data and indicated in the figure legends. P values were calculated using unpaired 2-tailed *t*-test and with Welch's correction when necessary, Mann-Whitney test, and one-way ANOVA followed by Dunnet's test. Paired t-test and Wilcoxon test was used to compare matched samples in Fig. **5J** and **5K**. P values of less than 0.05 were considered significant.

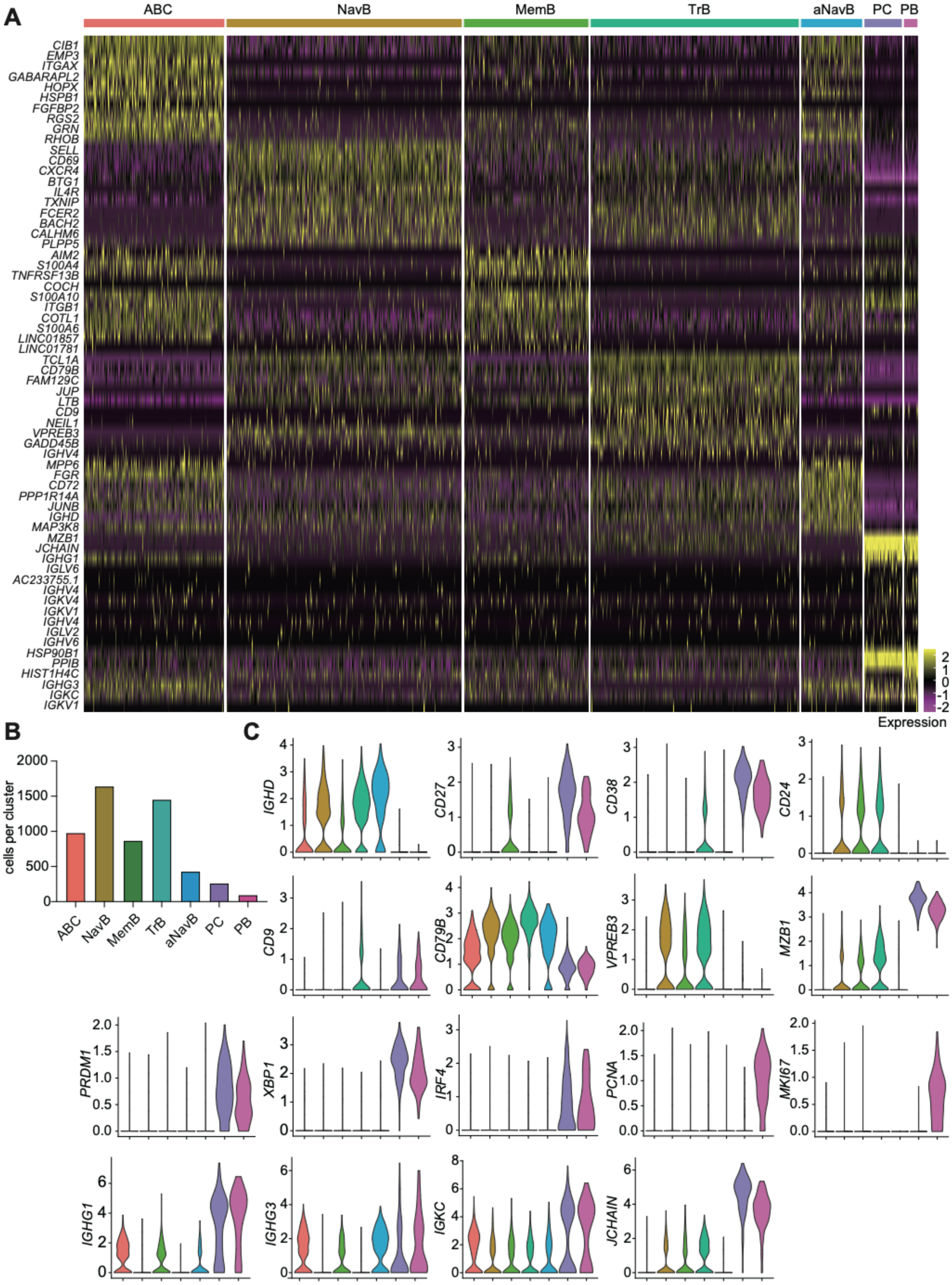
A
ABC
NavB
MemB
TrB
aNavB
PC
PB
CIB1
EMP3
ITGAX
GABARAPL2
HOPX
HSPB1
FGFBP2
RGS2
GRN
RHOB
SELL
CD69
CXCR4
BTG1
IL4R
TXNIP
FCER2
BACH2
CALHM6
PLPP5
AIM2
S100A4
TNFRSF13B
COCH
S100A10
ITGB1
COTL1
S100A6
LINC01857
LINC01781
TCL1A
CD79B
FAM129C
JUP
LTB
CD9
NEIL1
VPREB3
GADD45B
IGHV4
MPP6
FGR
CD72
PPP1R14A
JUNB
IGHD
MAP3K8
MZB1
JCHAIN
IGHG1
IGLV6
AC233755.1
IGHV4
IGKV4
IGKV1
IGHV4
IGLV2
IGHV6
HSP90B1
PPIB
HIST1H4C
IGHG3
IGKC
IGKV1
Expression
B
cells per cluster
ABC
NavB
MemB
TrB
aNavB
PC
PB
C
IGHD
CD27
CD38
CD24
CD9
CD79B
VPREB3
MZB1
PRDM1
XBP1
IRF4
PCNA
MKI67
IGHG1
IGHG3
IGKC
JCHAIN

**Fig. S1. B cells landscape from new-onset SLE patient.**

(**A**) Heat map representing scaled expression values of top 10 genes defining each cluster. (**B**) Number of cells per cluster. (**C**) Violin plots of select gene expression (*IGHD*, CD27, *CD38*, *CD24*, *CD9*, *CD79B*, *VPREB3*, *MZB1*, *PRDM1*, *XBP1*, *IRF4*, *PCNA*, *MKI67*, *IGHG1*, *IGHG3*, *IGKC*, *JCHAIN*) in different clusters, with log-normalized expression values labeled.

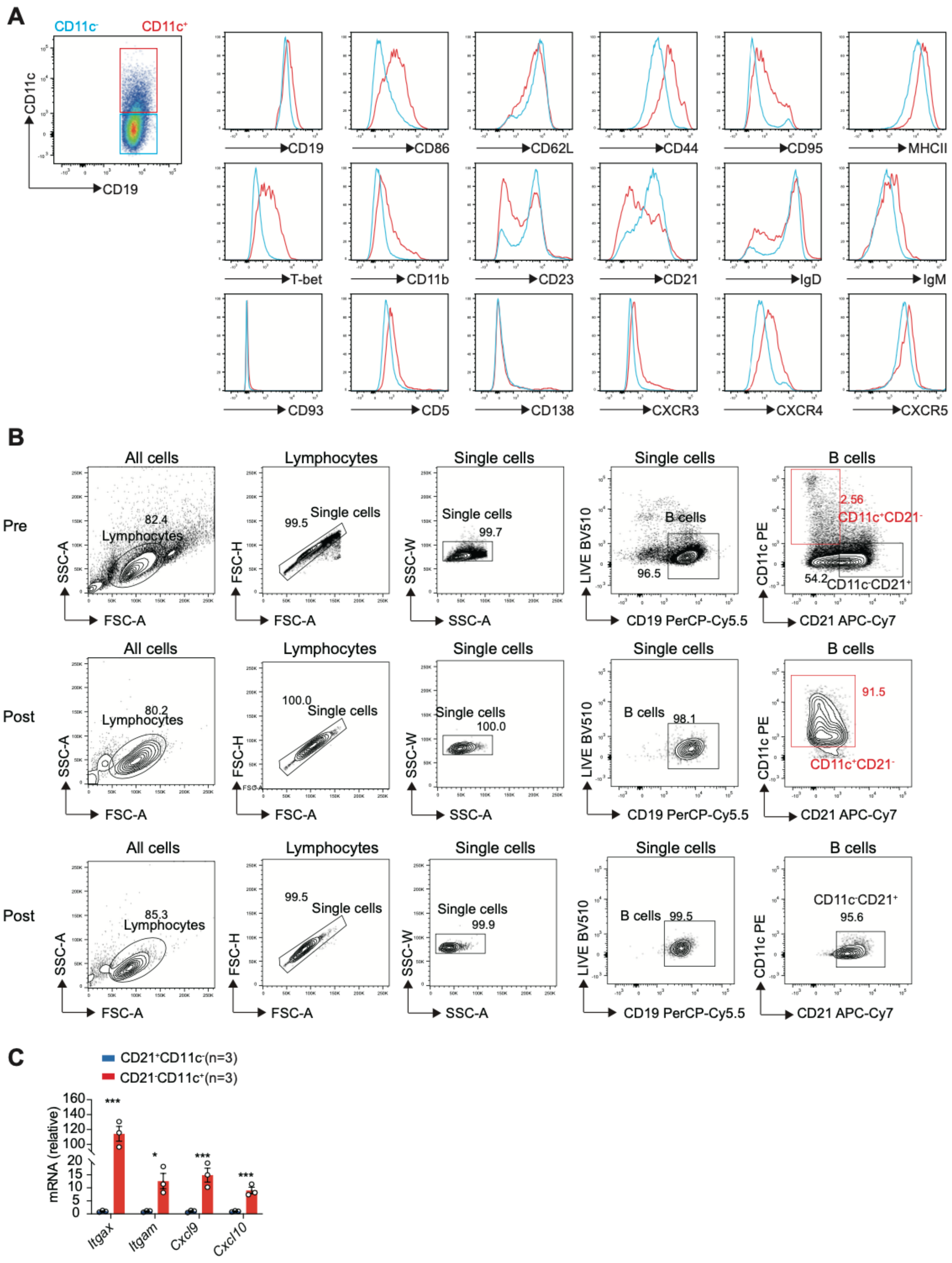
A
CD11c-
CD11c+
CD11c
CD19
CD86
CD62L
CD44
CD95
MHCII
T-bet
CD11b
CD23
CD21
IgD
IgM
CD93
CD5
CD138
CXCR3
CXCR4
CXCR5
B
Pre
Post
Post
All cells
Lymphocytes
Single cells
B cells
82.4
99.5
99.7
96.5
2.56
CD11c+CD21-
54.2
CD11c-CD21+
80.2
100.0
98.1
91.5
85.3
99.9
99.5
95.6
SSC-A
FSC-A
FSC-H
SSC-W
LIVE BV510
CD19 PerCP-Cy5.5
CD11c PE
CD21 APC-Cy7
C
CD21+CD11c-(n=3)
CD21-CD11c+(n=3)
mRNA (relative)
Itgax
Itgam
Cxcl9
Cxcl10

**Fig. S2. Phenotype of ABCs from cGVHD lupus mice.**

(**A**) C57/B6 mice were intraperitoneally injected with 10 million $CD4^+$ T cells from bm12 mice for 2 weeks and sacrificed for detection. Flow cytometry detection of splenic $CD11c^−$ (blue) or $CD11c^+$ (red) B cells (n=3) for characteristic markers. (**B**) Pre-and post-sort FACS gating strategy of splenic $CD19^+CD11c^+CD21^-$ and $CD19^+CD11c^-CD21^+$ B cells from mice described in **A**. (**C**) Realtime-PCR analysis of ABC signature genes (*Itgax*, *Itgam*, *Cxcl9*, *Cxcl10*) in splenic $CD19^+CD11c^+CD21^-$ and $CD19^+CD11c^-$ $CD21^+$ B cells (n=3) sorted by flow cytometry at week 2 after bm12 induction described in **A**. Data are representative of two in **A** and three in **B** and **C** independent experiments with 3 mice per group. Bars indicate mean ± SEM values. Statistical analysis was performed using unpaired t-test (**C**). *p<0.05, **P<0.01, ***P<0.001, ns, not significant.

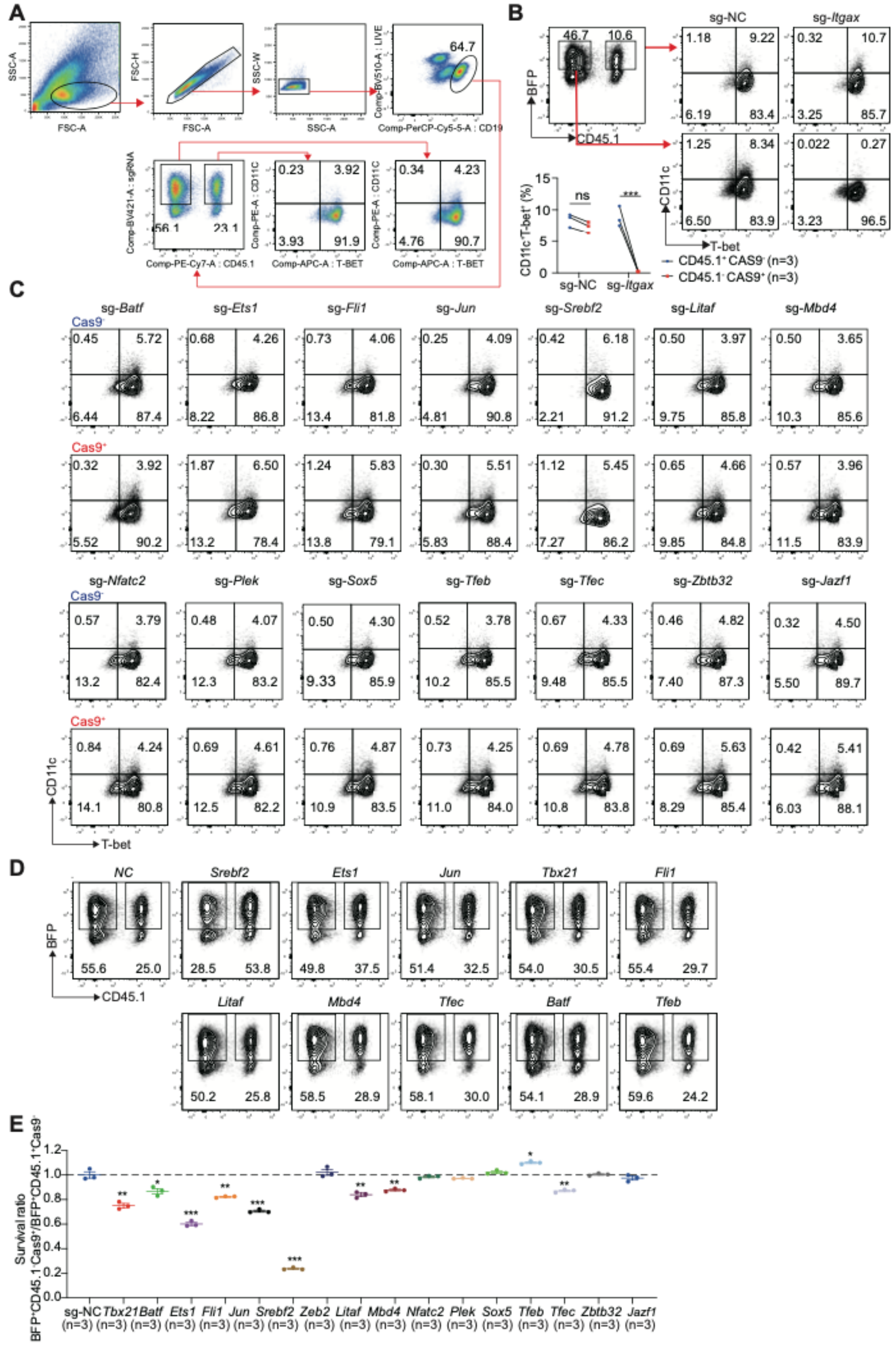

**Fig. S3. Gene editing in mouse primary B cells and screen TFs for ABC differentiation.**

(**A**) Splenic B cell were activated, edited, and induced for ABC differentiation described as in **Fig. 1E**. Gating strategy for flow cytometry analysis. Cells with certain size gated by FSC-A and SSC-A exclude duplicates by FSC-A/H and SSC-A/W. Then viable B cells were gated from CD19$^{+}$Zombie Aqua$^{-}$ and further divided into BFP$^{+}$CD45.1$^{-}$ fraction and BFP$^{+}$CD45.1$^{+}$ fraction. ABCs percentages were gated from CD11c$^{+}$T-bet$^{+}$ and were compared between BFP$^{+}$CD45.1$^{-}$

fraction and $BFP^+CD45.1^+$ fraction. The ABC percentage in $BFP^+CD45.1^-$ fraction versus percentage in $BFP^+CD45.1^+$ fraction was set and then sg-TF group was normalized with sg-NC group and calculated as ABC ratio. (**B**) Flow cytometry plots and frequency of ABCs ($CD11c^+$T-$bet^+$) in $CD45.1^+$sg-$Itgax^+$ ($Cas9^-$) and $CD45.1^-$sg-$Itgax^+$ ($Cas9^+$) cells described in **A**. (**C**) Flow cytometry plots of $CD11c^+$T-$bet^+$ B cells in $CD45.1^+sgRNA^+$ (Cas9-$sgRNA^+$) and $CD45.1^-$$sgRNA^+$ ($Cas9^+sgRNA^+$) cells targeting indicated genes. (**D**) Flow cytometry plots of $CD45.1^-$($GFP^+$) and $CD45.1^+$($GFP^-$) B cells targeting indicated genes after gating on $BFP^+$ cells. (**E**) Statistical analysis of cell survival ratio by comparing $CD45.1^-$($GFP^+$) with $CD45.1^+$($GFP^-$) B cells and then normalized with sg-NC. Data are representative of 4 independent experiments. Bars indicate mean ± SEM values. Statistical analysis was performed using paired t-test (**B**) and ordinary one-way ANOVA with two-sided Dunnett's multiple comparisons testing (**E**). *p<0.05, **P<0.01, ***P<0.001, ns, not significant.

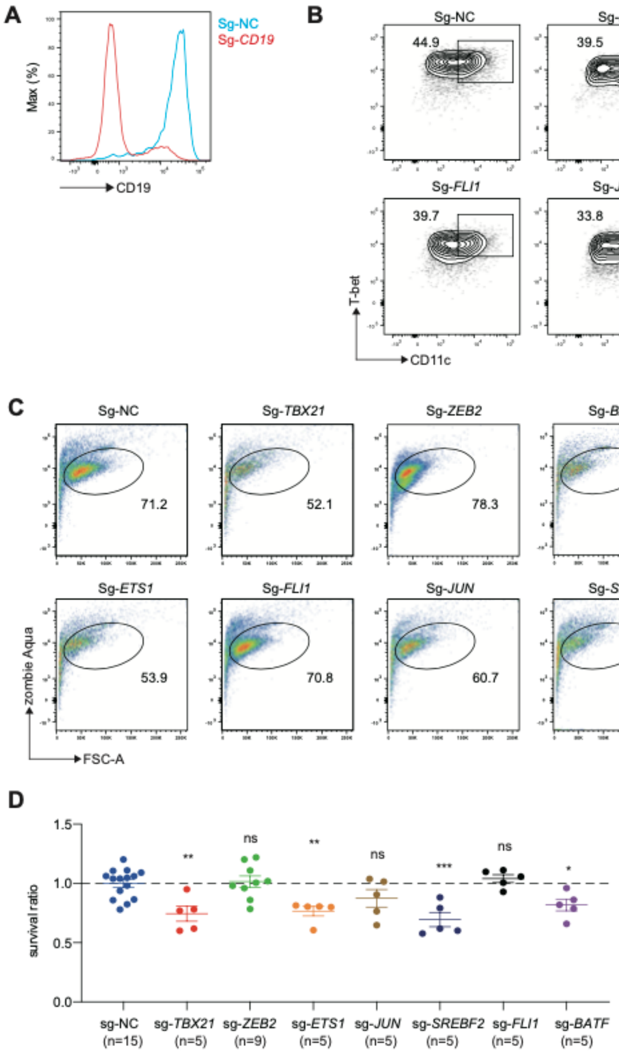


**Fig.S4. Screen TFs for human ABC differentiation.**
(**A**) Cas9 RNP delivery to primary B cells. Positive control Sg-*CD19* was used to verify the efficiency of gene editing in human primary B cells. (**B**) Flow cytometry plots of CD27$^-$IgD$^-$ CD11c$^+$T-bet$^+$ ABCs. Human B cells were activated and then were electroporated with Cas9-RNP targeting indicated gene. (**C and D**) Flow cytometry plots (**C**) and statistical analysis (**D**) of cell viability of B cells electroporated with Cas9-RNP targeting indicated gene and differentiated in ABC skewing condition. n refers to the measurement of distinct samples (biological repeats). Data are representative of 3 independent experiments. Bars indicate mean ± SEM values. Statistical analysis was performed using ordinary one-way ANOVA with two-sided Dunnett's multiple comparisons testing (**D**). *p<0.05, **P<0.01, ***P<0.001, ns, not significant.

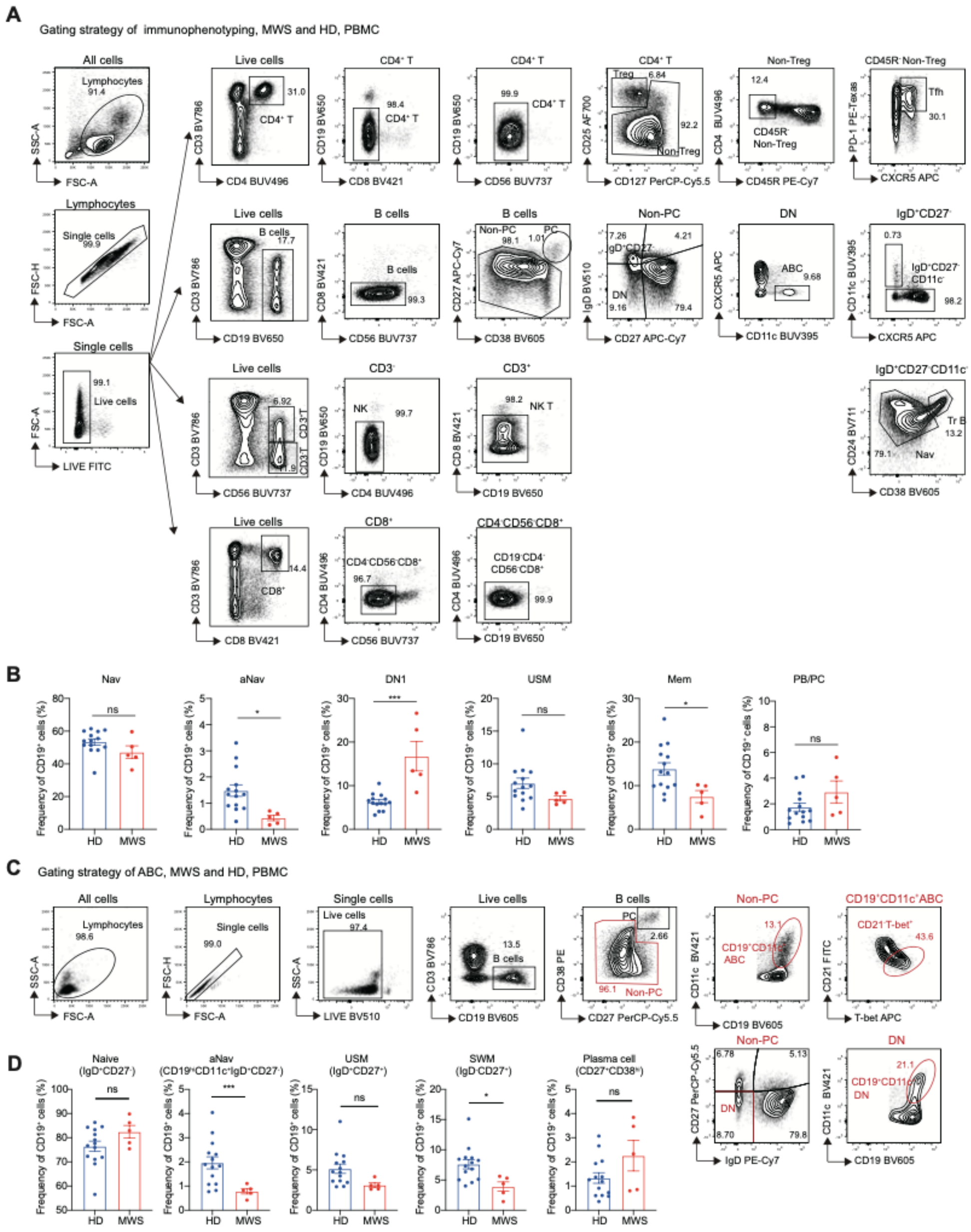

**Fig.S5. Immunological phenotype of MWS patients**

(**A**) Gating strategy for lymphocytes staining described in **Fig.2C**. (**B**) Statistical analysis of Nav, aNav, DN1, USM, Mem, PB/PC described in **Fig.2D** in PBMC for HD versus MWS group. (**C**) Gating strategy of ABCs in PBMC for HD and MWS group. (**D**) Statistical analysis of Naive, aNav, USM, SWM, plasma cells for HD versus MWS group described in (**C**). n refers to the measurement of distinct samples (biological repeats). Bars indicate mean ± SEM values. Statistical analysis was performed using unpaired t-test (**B**, Nav-Mem, **D,** Naïve, USM-Plasma cell) with Welch's correction (**D**, aNav) and Mann-Whitney test (**B**, PB/PC). *p<0.05, **P<0.01, ***P<0.001, ns, not significant.

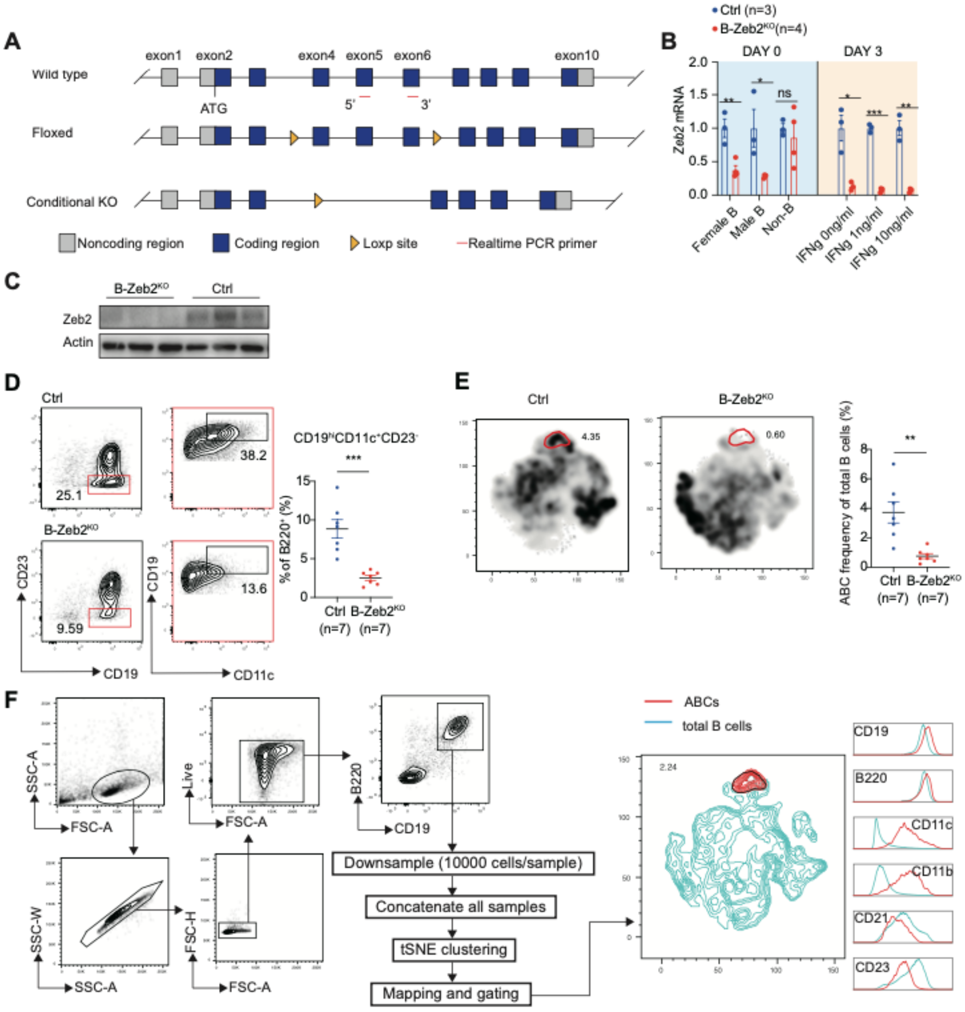


**Fig.S6. Construction of Zeb2 CKO mice and IMQ-induced lupus model.**
(**A**) Schematic diagram for construction of Zeb2 conditional knockout allele: Exon 4 to 6 are chosen as a target for inserting Loxp sites by using CRISPR/Cas9 technology. The floxed mice was knocked out after cross with mice expressing Cre recombinase. Realtime-PCR primers to detect Zeb2 expression are labeled in red. (**B**) Realtime-PCR analysis of Zeb2 expression in splenic B cells and non-B cell fractions (Day 0) or ABC skewing cocktail for 3 days. (**C**) Western blot analysis of Zeb2 protein level in splenic B cells from B-Zeb2$^{KO}$ and CD19$^{Cre/+}$ (Ctrl) mice. (**D**) Flow cytometry plots and statistical analysis of splenic ABCs (CD19$^{hi}$CD11c$^{+}$CD23$^{-}$) from B-Zeb2$^{KO}$ and CD19$^{Cre/+}$ (Ctrl) mice were induced by IMQ for 6 weeks described in **Fig. 3C**. (**E**) Separated t-SNE plots for two representative B-Zeb2$^{KO}$ (left) and Ctrl (right) samples. Statistical analysis of ABC in IMQ-induced B-Zeb2$^{KO}$ versus Ctrl mice. (**F**) Comprised t-SNE plots and gating strategy of splenic B cell clusters for IMQ-induced B-Zeb2$^{KO}$ and Ctrl mice. n refers to the measurement of distinct samples (biological repeats). Data are representative of 2 independent experiments. Bars indicate mean ± SEM values. Statistical analysis was performed using unpaired t-test (**B**, **D** and **E**). *p<0.05, **P<0.01, ***P<0.001, ns, not significant.

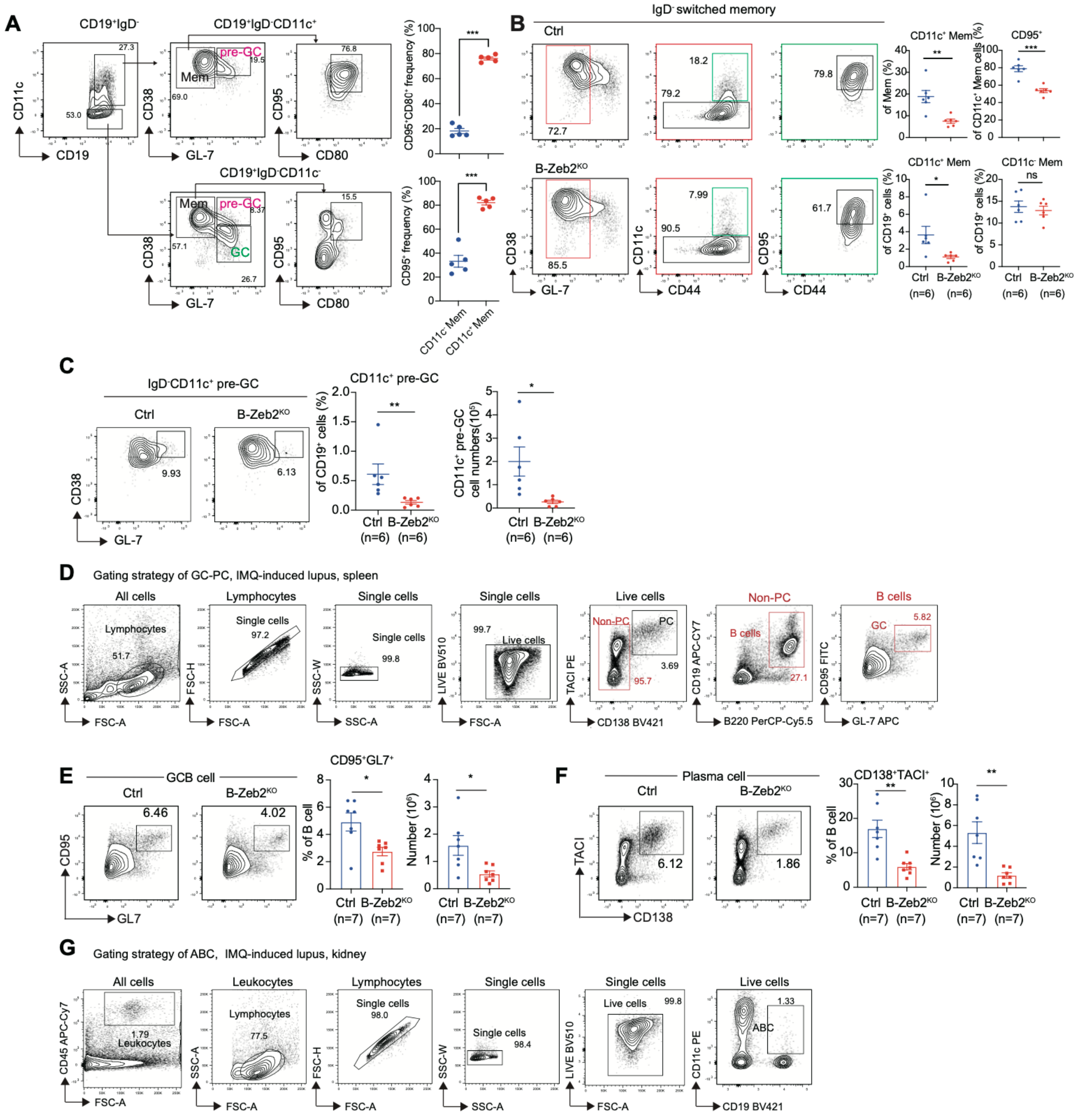


**Fig. S7. The effector B cell phenotype of IMQ-induced lupus in B cell conditional Zeb2 deficient mice**

(**A**) Gating strategy for the identification of the component of IgD$^-$CD19$^{hi}$CD11c$^+$ cells and IgD$^-$CD19$^{hi}$CD11c$^-$ cells. Statistical analysis of the percentage of CD95$^+$CD80$^+$ and CD95$^+$ in CD11c-/CD11c$^+$ memory B cells. (**B**) Gating strategy identifies CD11c$^+$ memory B cells, CD11c$^-$ memory B cells, CD44$^{hi}$CD95$^+$CD11c$^+$ memory B cells. Flow cytometry plots and statistical analysis of the above cell subsets from B-Zeb2$^{KO}$ mice and control mice described in **Fig. 3C**. (**C**) Flow cytometry plots and statistical analysis of CD11c$^+$ pre-GC-like B cells (CD19$^+$IgD$^-$CD11c$^+$CD38$^+$ GL-7$^+$) from B-Zeb2$^{KO}$ mice and control mice. (**D to F**) Gating strategy identifies splenic GCBs and Plasma cells**.** Flow cytometry plots and statistical analysis of the splenic GC B cells (CD19$^+$CD95$^+$GL-7$^+$, **E**), and plasma cells (CD19$^+$CD138$^+$TACI$^+$, **F**) from B-Zeb2$^{KO}$ mice and control mice described in **Fig. 3C**. (**G**) Gating strategy of ABC from kidney for **Fig.3J**. n refers to the measurement of distinct samples (biological repeats). Data are representative of 2 independent experiments. Bars indicate mean ± SEM values. Statistical analysis was performed using unpaired t-test (**A**, **B**, and **E** for frequency) with Welch's correction (**C** for number, **E** for number, and **F**) and Mann-Whitney test (**C** for frequency). *p<0.05, **P<0.01, ***P<0.001, ns, not significant.

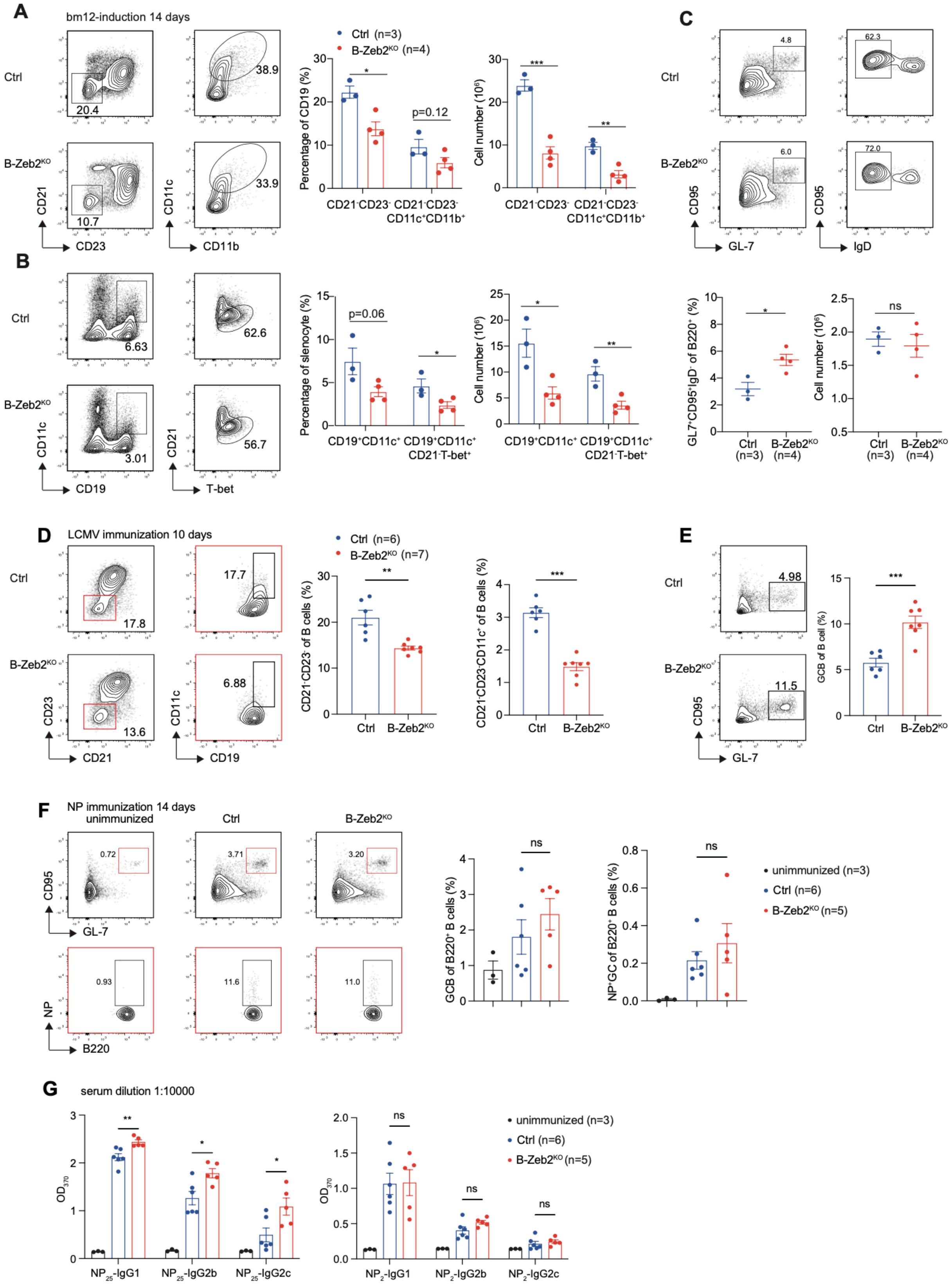
A
bm12-induction 14 days
Ctrl
B-Zeb2KO
20.4
10.7
38.9
33.9
CD21
CD23
CD11c
CD11b
Ctrl (n=3)
B-Zeb2KO (n=4)
Percentage of CD19 (%)
Cell number (10^6)
p=0.12
CD21-CD23-
CD21-CD23- CD11c+CD11b+
B
6.63
3.01
62.6
56.7
CD11c
CD19
CD21
T-bet
Percentage of slenocyte (%)
p=0.06
CD19+CD11c+
CD19+CD11c+ CD21-T-bet+
C
4.8
6.0
62.3
72.0
CD95
GL-7
IgD
GL7+CD95+IgD- of B220+ (%)
Cell number (10^6)
Ctrl (n=3)
B-Zeb2KO (n=4)
ns
D
LCMV immunization 10 days
17.8
13.6
17.7
6.88
CD23
CD21
CD11c
CD19
Ctrl (n=6)
B-Zeb2KO (n=7)
CD21-CD23- of B cells (%)
CD21-CD23-CD11c+ of B cells (%)
E
4.98
11.5
CD95
GL-7
GCB of B cell (%)
F
NP immunization 14 days
unimmunized
Ctrl
B-Zeb2KO
0.72
3.71
3.20
0.93
11.6
11.0
CD95
GL-7
NP
B220
GCB of B220+ B cells (%)
NP+GC of B220+ B cells (%)
unimmunized (n=3)
Ctrl (n=6)
B-Zeb2KO (n=5)
G
serum dilution 1:10000
OD370
NP25-IgG1
NP25-IgG2b
NP25-IgG2c
NP2-IgG1
NP2-IgG2b
NP2-IgG2c
unimmunized (n=3)
Ctrl (n=6)
B-Zeb2KO (n=5)

**Fig. S8. The effector B cell phenotype in different immunization models.**
(**A** to **C**) B-Zeb2$^{KO}$ mice and control mice were intraperitoneally injected with 10 million CD4$^{+}$ T cells from bm12 mice for 14 days and sacrificed for detection. Flow cytometry plots and statistical analysis of the percentage and numbers of splenic ABCs (CD21$^{-}$CD23$^{-}$ B cells, CD21$^{-}$CD23$^{-}$ CD11c$^{+}$CD11b$^{+}$ B cells, **A**, and CD19$^{+}$CD11c$^{+}$, CD19$^{+}$CD11c$^{+}$CD21$^{-}$T-bet$^{+}$, **B**) and GCB cells (IgD$^{-}$GL-7$^{+}$CD95$^{+}$, **C**) from B-Zeb2$^{KO}$ mice and control mice as described above. (**D**-**E**) The effector B cell phenotype in LCMV immunization model. B-Zeb2$^{KO}$ mice and control mice were infected with LCMV for 10 days. Representative flow cytometry plots and statistical analysis of CD21$^{-}$CD23$^{-}$ B cells, CD21$^{-}$CD23$^{-}$ CD11c$^{+}$ ABCs (**D**) and GCBs (GL-7$^{+}$CD95$^{+}$, **E**) from B-Zeb2$^{KO}$ mice and control mice. (**F-G**) B-Zeb2$^{KO}$ mice and control mice were immunized with NP-CGG (100ug/mice) plus alum and LPS (1ug/mice) adjuvants for 14 days. (**F**) Representative flow cytometry plots and statistical analysis of GCB cells (GL-7$^{+}$CD95$^{+}$) and NP-specific GCB cells (NP$^{+}$GL-7$^{+}$CD95$^{+}$) from B-Zeb2$^{KO}$ mice, control mice and unimmunized mice. (**G**) $NP_{25}$-specific and $NP_{2}$-specific IgG1, IgG2b and IgG2c antibodies were detected in the serum from B-Zeb2$^{KO}$ mice and control mice. n refers to the measurement of distinct samples (biological repeats). Data are representative of 2 independent experiments. Bars indicate mean ± SEM values. Statistical analysis was performed using unpaired t-test (**A** to **G**). *p<0.05, **P<0.01, ***P<0.001, ns, not significant.

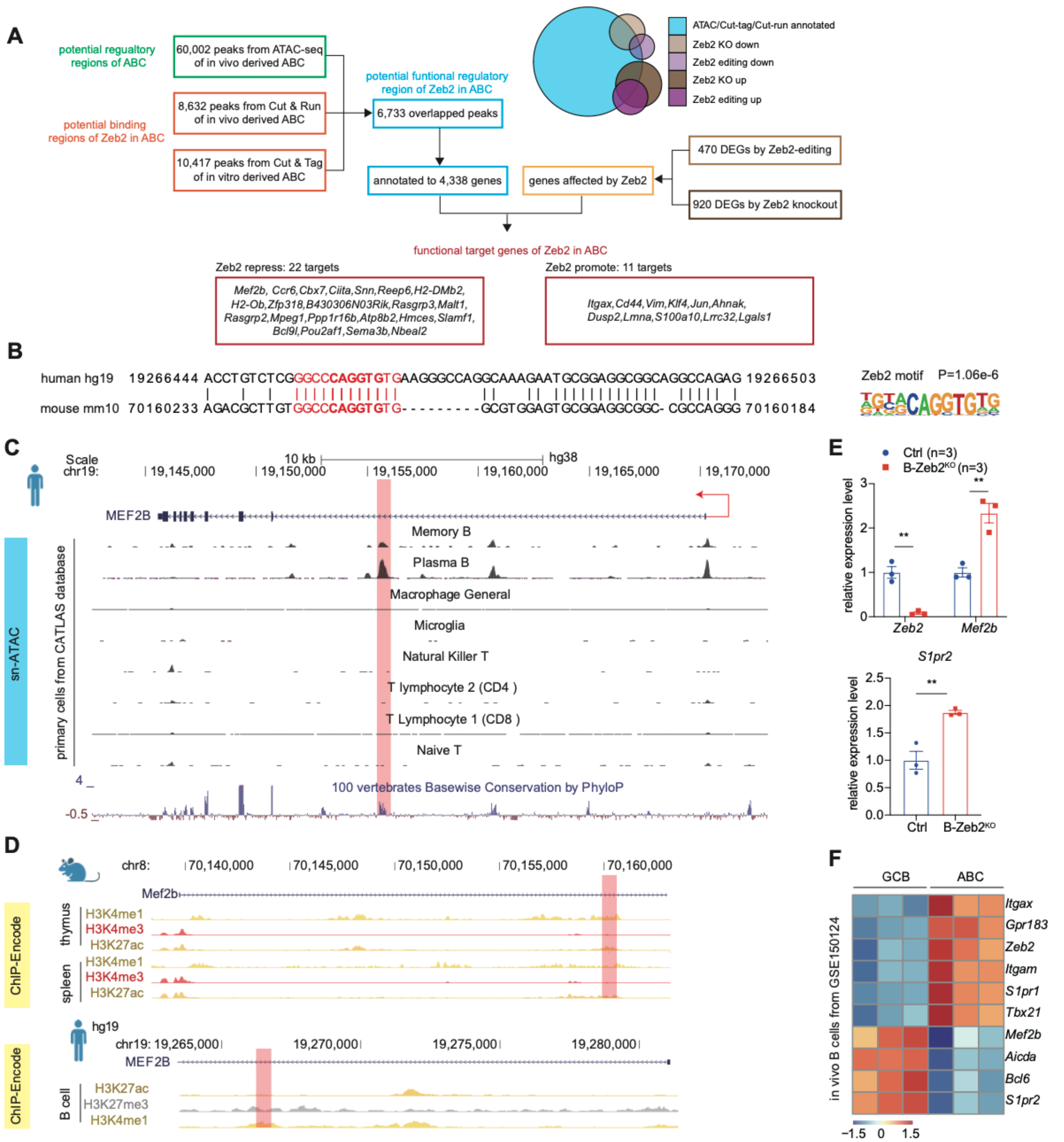


**Fig. S9. Zeb2 directly binds to the intronic enhancer of Mef2b to interfere its expression**

(**A**) The diagram of analysis strategy for screening and identifying functional target genes of Zeb2 in ABC. The chromatin accessibility (peaks from ATAC) of ABCs was compared with two transcription factor binding site characterization approaches to reveal Zeb2 binding sites: CUT & Tag and CUT & RUN. After overlapping, 6,733 accessible sites with Zeb2 binding were identified and annotated to 4,338 genes. Among the genes differentially expressed by Zeb2 deficiency, 33 candidate target genes of Zeb2 were identified, with 22 genes repressed and 11 genes activated by Zeb2. (**B**) The conserved sequence of human and mouse in Mef2b +20kb intronic enhancer region with a Zeb2 binding motif. (**C**) The chromatin accessibility around Mef2b locus in human primary immune cell subsets from single nuclear ATAC data (CATLAS database) mapped in hg18. (**D**) ChIP-seq tracks display histone modifications around the Mef2b locus in human (hg19) and mouse (mm10) from Encode database. (**E**) Realtime-PCR validation of Zeb2, Mef2b and S1pr2 expression between Zeb2-deficient B cells and Ctrl B cells. (**F**) The heatmap shows the expression of the selected genes between ABC and GCB from GSE150124. n refers to the measurement of distinct samples (biological repeats). Data are representative of 2 independent experiments (**E**). Bars indicate mean ± SEM values. Statistical analysis was performed using unpaired t-test (**E**). *$p<0.05$, **$P<0.01$, ***$P<0.001$, ns, not significant.

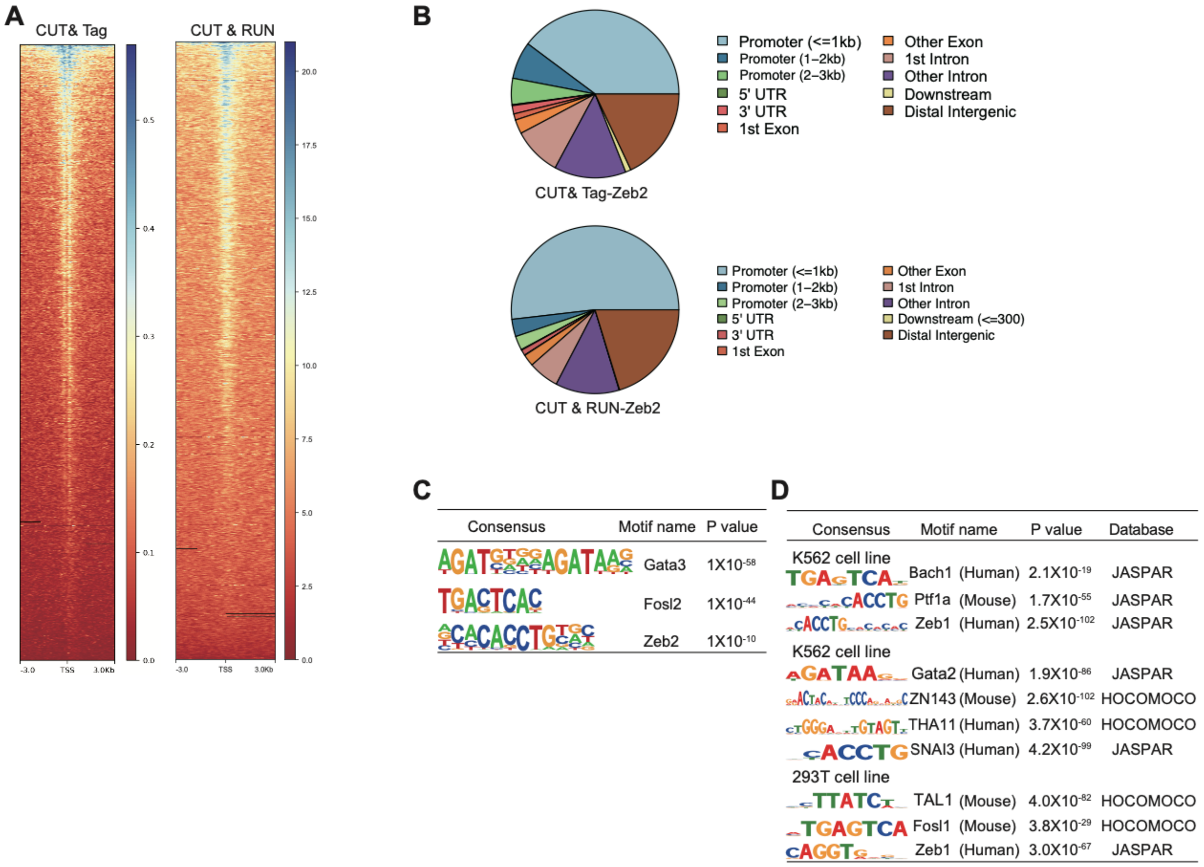


**Fig. S10. The transcriptional regulation of Zeb2 in mouse ABC was analyzed by CUT&Tag and CUT & RUN.**

(**A**) Heatmap graph showing distributions of Zeb2 specific signaling from CUT&Tag and CUT & RUN. The horizontal axis represented the normalized gene range coordinates. (**B**) Peak annotation analysis of distribution of Zeb2 in the functional area of gene covered. (**C**) Motif-enrichment analysis of Zeb2 specific peaks generated from CUT&Tag sequencing data of in vitro derived ABCs. (**D**) Motif analysis of Zeb2 specific peak of Chip-seq data in cell lines from Encode project.

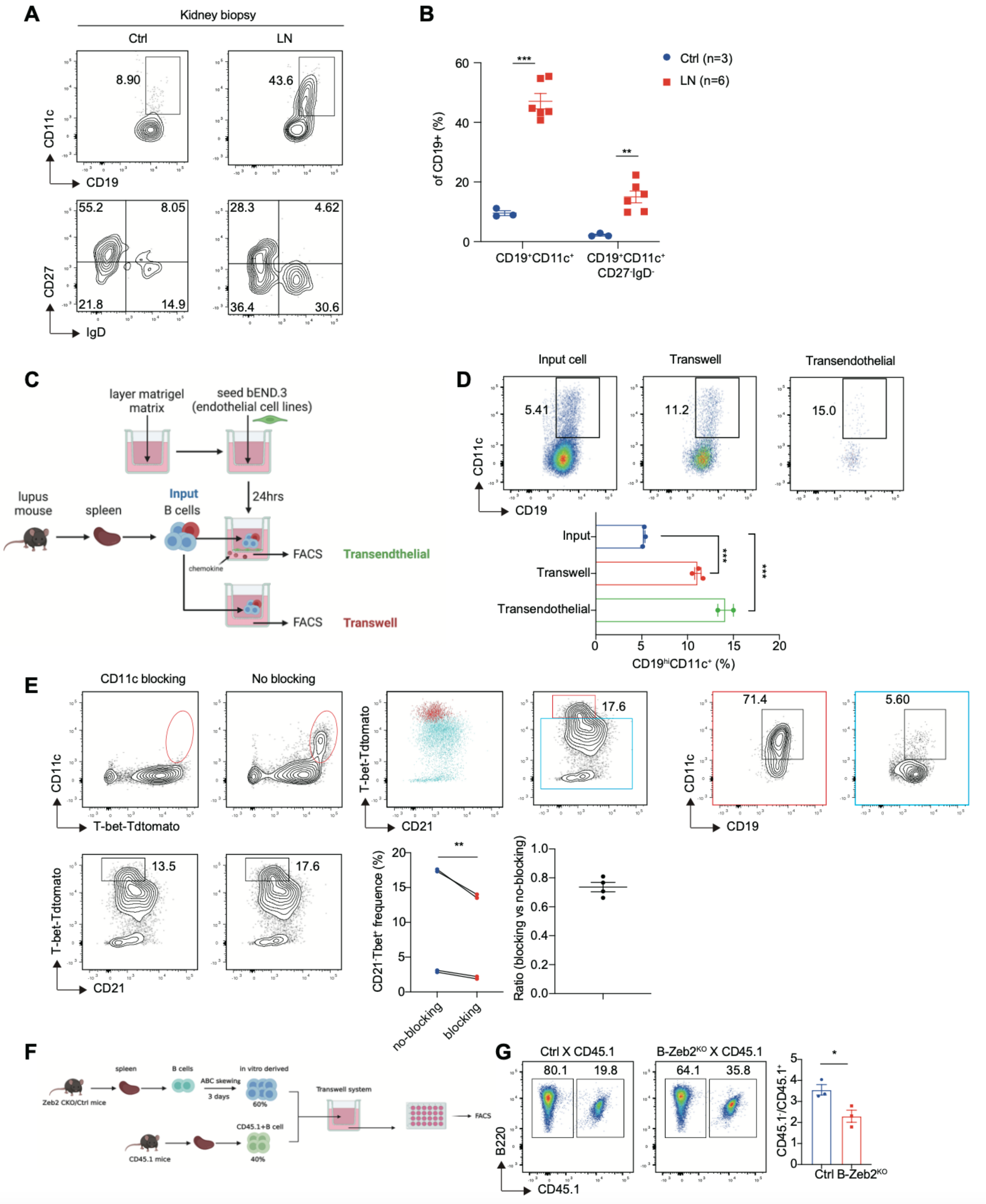


**Fig. S11. ABCs acquire enhanced migratory capacity via upregulated expression of beta2 integrin CD11c controlled by Zeb2.**

(**A-B**) Representative plots and statistical analysis of renal $CD19^+CD11c^+$ B cells and $CD19^+CD11c^+CD27^-IgD^-$ B cells from para-cancer control (Ctrl) and lupus nephritis (LN). (**C**) The diagram of trans-well and trans-endothelial system for migration assay. (**D**) Representative plots and frequency of the $CD19^{hi}CD11c^{+}$ among input control, trans-well and trans-endothelial groups. (**E**) Representative plots and frequency of the migrated ABC with or without CD11c blocking. (**F**) Flow chart of in vitro mouse ABC migration assay by trans-well system. B cells were isolated from B-Zeb2$^{KO}$ mice and control mice spleen and cultured under ABC-skewing condition for three days. (**G**) Representative plots and statistical analysis of the migratory capacity of in vitro derived mouse ABC in (**F**). n refers to the measurement of distinct samples (biological repeats). Data are representative of 2 independent experiments. Bars indicate mean ± SEM values. Statistical analysis was performed using unpaired t-test (**B** and **G**) with Welch's correction (**B**, LN vs. Ctrl of $CD19^{+}CD11c^{+}CD27^{-}IgD^{-}$), ordinary one-way ANOVA with two-sided Dunnett's multiple comparisons testing (**D**), and paired t-test (**E**). *p<0.05, **P<0.01, ***P<0.001, ns, not significant.

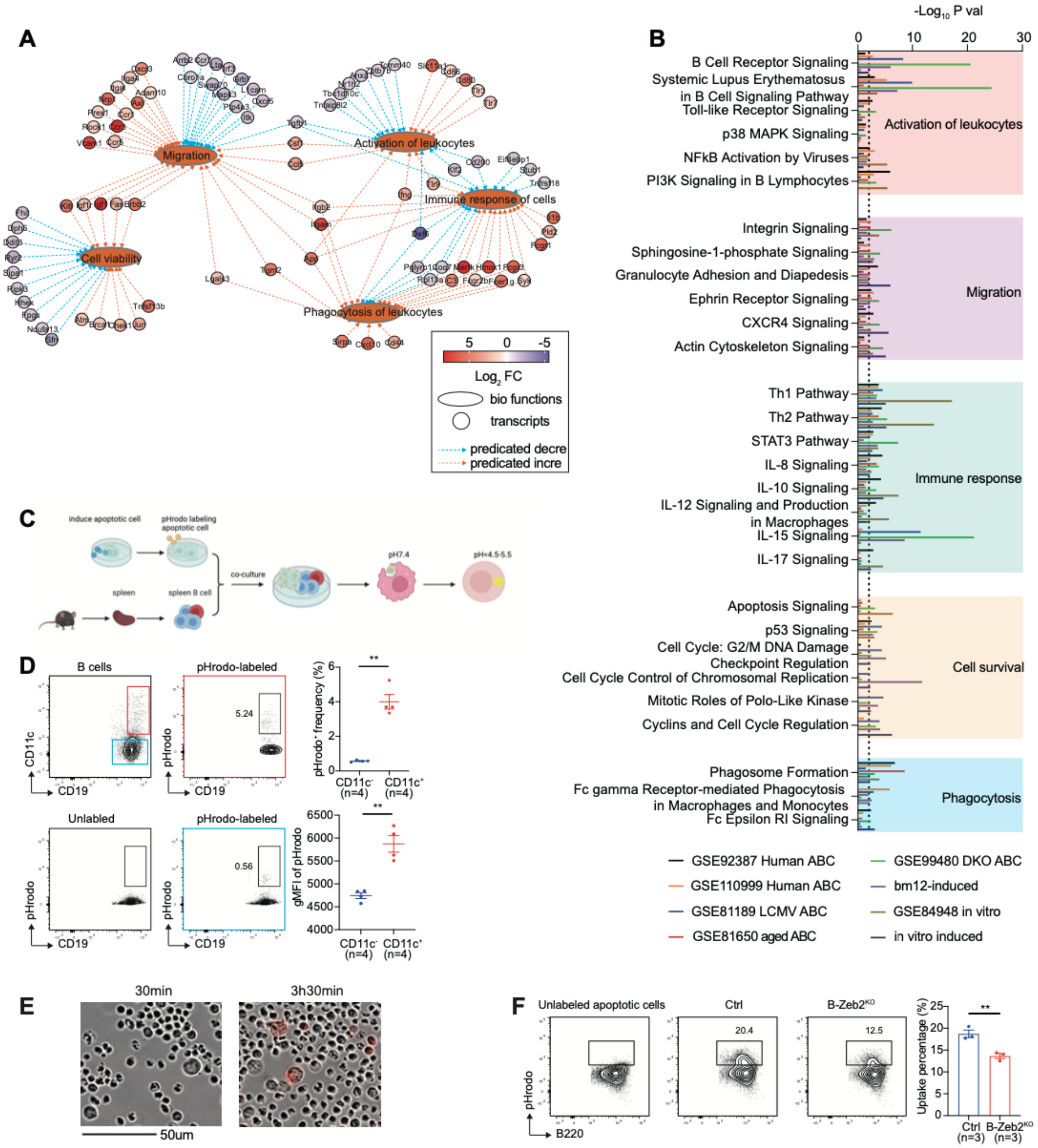


**Fig. S12. The distinct phagocytic feature of ABCs was regulated by Zeb2**

(**A**) Biological functional classification and network analysis of differentially-expressed genes (DEGs) using Ingenuity Pathways Analysis (IPA). DEGs were overlaid to the network to find a biological regulatory effect based on previously reported interactions in the literature. The dotted line indicates the predicted relationship between genes and biological function: orange representing activation, blue representing inhibition. Cell activation, viability, migration, immune response and phagocytosis pathway are activated in ABCs. (**B**) Pathway enrichment analysis of ABC from different datasets described in **Fig. 4, B** and **G**, The charts represent the top significantly signaling pathways related with functional categories: activation of leukocytes, migration, immune response, cell survival and phagocytosis. (**C**) Flow chart of the in vitro phagocytosis assay. (**D**) Representative plots and statistical analysis of the pHrodo-labeled apoptotic cells uptake percentage and intensity by splenic $CD11c^+$ and $CD11c^-$ B cells from lupus mice. (**E**) Representative microscopy images of phagocytosis of pHrodo-labeled apoptotic cells by ABC at indicated time points. Scale bar = 50 μm. (**F**) B cells isolated from B-$Zeb2^{KO}$ mice and control mice spleen and cultured under ABC-skewing condition for three days. Representative plots and statistical analysis of the pHrodo-labeled apoptotic cells uptake percentage by in vitro derived B cells. n refers to the measurement of distinct samples (biological repeats). Data are representative of 2 independent experiments. Bars indicate mean ± SEM values. Statistical analysis was performed using unpaired t-test (**D** and **F**). *p<0.05, **P<0.01, ***P<0.001, ns, not significant.

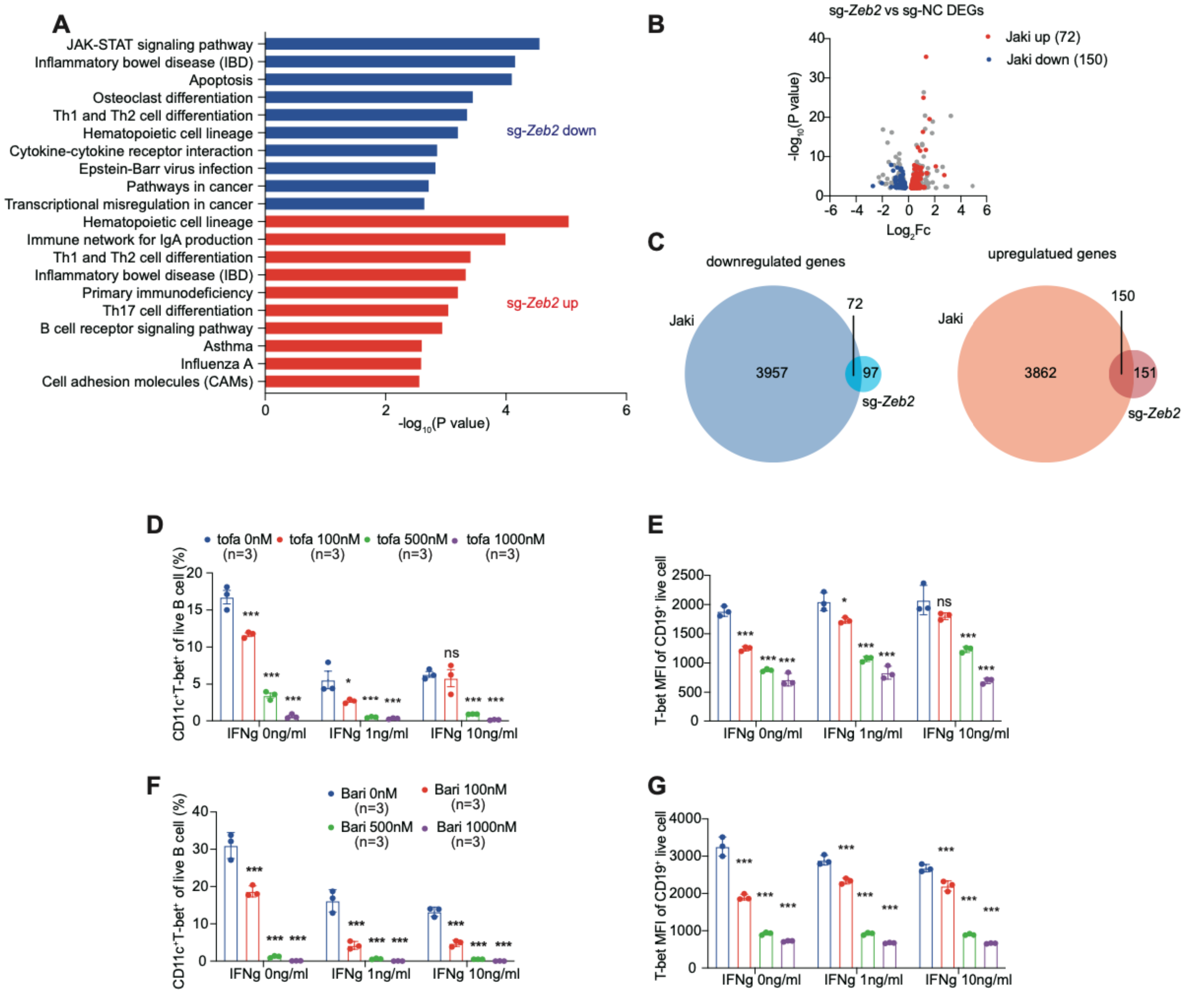


**Fig.S13. Jak-stat inhibitors impair in vitro derived ABCs formation.**

(**A**) KEGG pathway enrichment analysis of downregulated genes (blue) and upregulated genes (red) in sg-*Zeb2* B cells was performed by online Enrichr tools. P-values was log-normalized. (**B**) The Volcano plot shows the up-/down-regulated genes of Jaki among the DEGs determined by sg-*Zeb2* editing. (**C**) Venn diagram illustrating the overlap of up-/down-regulated genes between the sg-*Zeb2* editing and Jaki treated. (**D to G**) Statistical analysis of in vitro induced mouse CD11c$^{+}$T-bet$^{+}$ ABCs in addition of different concentration (mock, 100 nM, 500 nM and 1000 nM) of tofacitinib (**D**) or baricitinib (**F**). T-bet MFI of in vitro induced mouse CD11c$^{+}$T-bet$^{+}$ ABCs in addition of different concentration (mock, 100 nM, 500 nM and 1000 nM) of tofacitinib (**E**) or baricitinib (**G**). n refers to the measurement of distinct samples (biological repeats). Data are representative of 2 independent experiments. Bars indicate mean ± SEM values. Statistical analysis was performed using ordinary one-way ANOVA with two-sided Dunnett's multiple comparisons testing (**D** to **G**). *p<0.05, **P<0.01, ***P<0.001, ns, not significant.

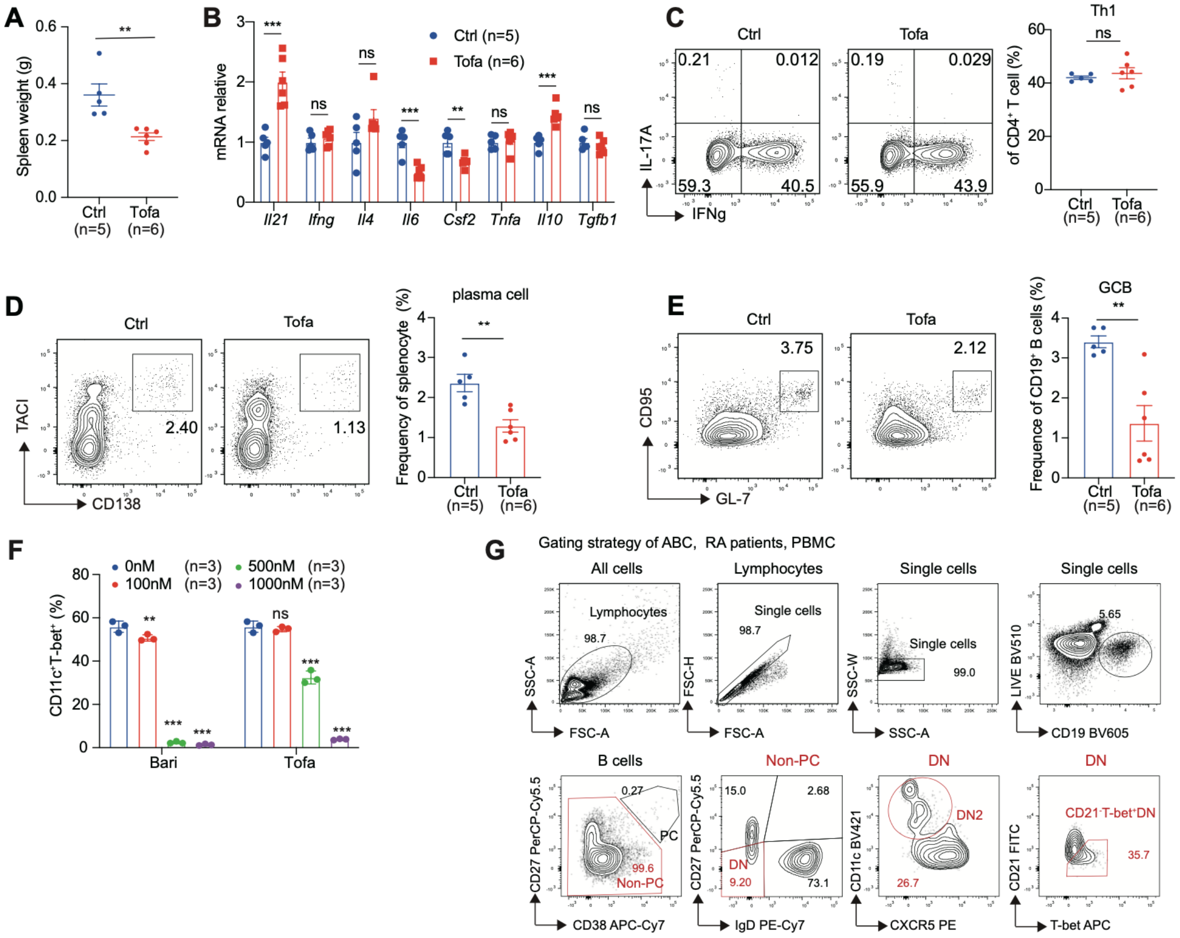


**Fig.S14 Jak-stat inhibitors impair in vivo derived ABCs formation.**

(**A**) The bm12 induced mice were treated with tofacitinib 30 mg/kg once a day by oral gavage for 2 weeks and then sacrificed. Changes of spleen weight in mice with tofacitinib treatment. (**B**) The expression of proinflammatory cytokines in splenocytes from Tofa-treated and control mice by qPCR. (**C** to **E**) Flow cytometry plots and statistical analysis of the Th1 (**C**), plasma cells (**D**), and GCB cells (**E**) from Tofa-treated and control mice described in (**A**). (**F**) Statistical analysis of in vitro induced human $CD11c^{+}T\text{-}bet^{+}$ ABCs in addition of different concentration (mock, 100 nM, 500 nM and 1000 nM) of tofacitinib or baricitinib. (**G**) Gating strategy of ABCs from PBMC for RA patients with tofacitinib treatment. Data are representative of 2 or 3 independent experiments in b-g. Bars indicate mean ± SEM values. Statistical analysis was performed using unpaired t test (**A** to **E**) and ordinary one-way ANOVA with two-sided Dunnett's multiple comparisons testing (**F**). *p<0.05, **P<0.01, ***P<0.001, ns, not significant.

| | SLE patient |
|---|---|
| Age | 27 |
| Sex | Female |
| WBC ($\times 10^9$/L) | 4.35 |
| RBC ($\times 10^{12}$/L) | 3.14 |
| Hb(g/L) | 106 |
| PLT($\times 10^9$/L) | 212 |
| ESR(mm/h) | 82 |
| IgA(g/L) | 4.23 |
| IgM(g/L) | 1.5 |
| IgG(g/L) | 26.5 |
| C3(g/L) | 0.149 |
| C4(g/L) | <0.017 |
| Anti-dsDNA(IU/ml) | 292.16 |
| SLEDAI | 24 |
| ANA | Positive |
| Anti-SSA | Negative |
| Anti-SSB | Negative |
| Anti-U1RNP | Negative |
| Anti-Sm | Negative |
| Anti-rRNP | Positive |
| Anti-ANUA | Positive |
| ACL | Positive |
| Fever | No |
| Rash | Yes |
| Alpecia | Yes |
| Arthritis | No |
| Oral ulcers | No |
| Cutaneous vasculitis | No |
| Raynaud's | No |
| Leukopenia and/or Thrombocytopenia | No |
| Anemia | No |
| Serositis | No |
| NPSLE | No |
| LN | III+V |
| Urine protein of LN patients (mg/24h) | 3379.8 |

**Table S1. Basic clinical characteristics of a patient with new-onset SLE**

| | LN biopsy(n=6) |
|---|---|
| Age (years),mean (SD) | 31.5 (5.32) |
| Female, n (%) | 6 (100) |
| SLEDAI | 10.5 (2.2) |
| rSLEDAI | 4 (0) |
| 24h upro (g/24h) | 4.3 (3.0) |
| ESR (mm/h) | 33.0 (24.9) |
| IgG (g/L) | 12.9 (9.1) |
| IgA (g/L) | 2.9 (0.7) |
| IgM (g/L) | 0.8 (0.4) |
| C3 (g/L) | 0.4 (0.1) |
| C4 (g/L) | 0.08 (0.05) |
| dsDNA (IU/ml) | 63.9 (30.6) |
| eGFR-EPI | 88.3 (18.6) |
| ANA | 1067 (330) |
| Renal pathology (ISN/RPS classification), n (%) | |
| II | 0 (0) |
| III | 0 (0) |
| IV | 3 (50) |
| V | 0 (0) |
| III+V | 0 (0) |
| IV+V | 3 (50) |

*SD:Standard deviation; n:number

**Table S2. Basic clinical characteristics of patients with lupus nephritis**

| RA patients | Before (n=6) Median±IQR | After (n=6) Median±IQR |
|---|---|---|
| WBC(× $10^9$/L) | 6.30±2.92 | 6.30±2.03 |
| RBC(× $10^{12}$/L) | 4.73±0.56 | 4.70±0.62 |
| Hb(g/L) | 136±25.25 | 137±23.50 |
| PLT(× $10^9$/L) | 322±127.5 | 274±120 |
| ESR(mm/h) | 28±25.88 | 19±11.00 |
| CRP(mg/L) | 10.4±32.80 | 6.6±4.90 |
| Cr(μmol/L) | 56.2±8.93 | 62.2±13.85 |
| ALT(U/L) | 21±11.25 | 25±7.00 |

**Table S3. Clinical characteristics of RA patients treatment with tofacitinib**

| Patient ID | Sex | age(year) | Variants in *ZEB2* | dbSNP ID | Inheritance |
|---|---|---|---|---|---|
| MWS-A1 | M | 5 | c.1027C>T<br>p.Arg343X | rs786204815 | *De novo* |
| MWS-B1 | F | 2 | c.2851C>T<br>p.Gln951X | - | *De novo* |
| MWS-C1 | M | 4 | c.1005delT<br>p.Ile335Metfs*2 | - | *De novo* |
| MWS-D1 | F | 4 | chr2:138434153-145285163 del<br>*ZEB2* loss exon 1-10 | - | *De novo* |
| MWS-E1 | F | 1 | chr2:145147017-145274917 del<br>*ZEB2* loss exon 2-10 | - | *De novo* |

**Table S4. Demographic and genetic information of MWS patients**

| Patient ID | Symptoms | | | | | | Infectious disease | Autoimmune disease | |
|---|---|---|---|---|---|---|---|---|---|
| | Distinctive facial features | Developmental delay | Intellectual disability | Epilepsy | Hirschsprung disease | Congenital malformations | upper respiratory tract infection (URTI) | ANA (1:80) | ENA[1] (1:100) |
| MWS-A1 | + | + | + | + | + | + | 4-5 times a year | - | - |
| MWS-B1 | + | + | + | + | - | + | 12 times a year | - | - |
| MWS-C1 | + | + | + | - | - | - | 3-4 times a year | - | - |
| MWS-D1 | + | + | + | + | - | - | 3 times a year | - | - |
| MWS-E1 | + | + | + | + | - | + | - | - | - |

+ feature present; - feature absent

[1] including anti-nRNP/Sm, anti-Sm, anti-SS-A, anti-Ro-52, anti-SS-B, anti-Scl-70, anti-Jo-1 and Anti-Rib-P

**Table S5. MWS patients clinical features**

| Biological Functions Annotation | P-value | Predicted Activation State | Activation z-score |
|---|---|---|---|
| Cell viability | 3.24E-24 | Increased | 6.955 |
| Cell survival | 3.04E-25 | Increased | 6.793 |
| Cell movement | 1.15E-33 | Increased | 6.462 |
| Migration of cells | 4.42E-31 | Increased | 6.432 |
| Phagocytosis | 6.71E-20 | Increased | 5.94 |
| Internalization of cells | 3.95E-14 | Increased | 5.919 |
| Phagocytosis of cells | 1.37E-19 | Increased | 5.878 |
| Engulfment of cells | 7.49E-20 | Increased | 5.775 |
| Organization of cytoskeleton | 4.15E-32 | Increased | 5.733 |
| Organization of cytoplasm | 9.07E-34 | Increased | 5.677 |
| Immune response of cells | 5.3E-27 | Increased | 5.577 |
| Homing of cells | 1.62E-16 | Increased | 5.406 |
| Engulfment of phagocytes | 2.64E-17 | Increased | 5.355 |
| Engulfment of blood cells | 2.36E-14 | Increased | 5.333 |
| Engulfment of myeloid cells | 9.02E-16 | Increased | 5.332 |
| Interaction of blood cells | 2.15E-20 | Increased | 5.328 |
| Engulfment of leukocytes | 2.6E-17 | Increased | 5.305 |
| Binding of blood cells | 2.07E-20 | Increased | 5.249 |
| Endocytosis | 3.33E-18 | Increased | 5.245 |
| Endocytosis by eukaryotic cells | 2.98E-15 | Increased | 5.225 |
| Microtubule dynamics | 8.08E-26 | Increased | 5.215 |
| Chemotaxis | 2.79E-16 | Increased | 5.174 |
| Engulfment by macrophages | 4.51E-15 | Increased | 5.14 |
| Adhesion of blood cells | 1.48E-20 | Increased | 5.017 |
| Engulfment of antigen presenting cells | 5.29E-15 | Increased | 5.015 |
| Binding of leukocytes | 9.89E-21 | Increased | 5.01 |
| Response of myeloid cells | 2.18E-20 | Increased | 5 |
| Phagocytosis of myeloid cells | 1.86E-16 | Increased | 4.938 |
| Phagocytosis of blood cells | 1.78E-13 | Increased | 4.857 |
| Response of phagocytes | 1.29E-21 | Increased | 4.854 |
| Phagocytosis of phagocytes | 3.17E-16 | Increased | 4.845 |
| Phagocytosis of leukocytes | 6.94E-17 | Increased | 4.819 |
| Immune response of myeloid cells | 2.44E-17 | Increased | 4.809 |
| Activation of cells | 1.65E-34 | Increased | 4.779 |

**Table S6. Top 35 biological function analysis of DSE99480 DKO ABC vs WT FoB**

| Antibodies | Source | Identifier |
|---|---|---|
| anti-mouse CD4 | biolegend | Clone GK1.5 |
| anti-mouse CD8 | biolegend | Clone 53-6.7 |
| anti-mouse CD19 | biolegend | Clone 6D5 |
| anti-mouse CD19 | biolegend | Clone 1D3/CD19 |
| anti-mouse CD44 | BD bioscience | Clone IM7 |
| anti-mouse CD62L | BD bioscience | Clone MEL-14 |
| anti-mouse PD-1 | eBioscience | Clone J43 |
| anti-mouse B220 | biolegend | Clone RA3-6B2 |
| anti-mouse CD95 | BD bioscience | Clone Jo2 |
| anti-mouse GL-7 | BD bioscience | Clone GL7 |
| anti-mouse IgM | BD Bioscience | Clone R6-60.2 |
| anti-mouse IgD | BD Bioscience | Clone 11-26c.2a |
| anti-mouse CD138 | biolegend | Clone 281-2 |
| anti-mouse TACI | biolegend | Clone 8F10 |
| anti-mouse CD11c | biolegend | Clone N418 |
| anti-mouse/human T-bet | biolegend | Clone 4B10 |
| anti-mouse CD21 | biolegend | Clone 7E9 |
| anti-mouse CD73 | biolegend | Clone TY/11.8 |
| anti-mouse CD80 | biolegend | Clone 16-10A1 |
| anti-mouse CD93 | BD Bioscience | Clone AA4.1 |
| anti-mouse CD45.1 | biolegend | Clone A20 |
| anti-mouse CD86 | BD bioscience | Clone GL1 |
| anti-mouse I-A/I-E | BD bioscience | Clone M5/114.15.2 |
| anti-mouse CD23 | BD bioscience | Clone B3B4 |
| anti-mouse CD11b | biolegend | Clone M1/70 |
| anti-mouse CD5 | eBioscience | Clone 53-7.3 |
| anti-mouse CXCR4 | BD Bioscience | Clone 2B11/CXCR4 |
| anti-mouse CXCR3 | biolegend | Clone CXCR3-173 |
| anti-human CD19 | biolegend | Clone HIB19 |
| anti-human CXCR5 | biolegend | Clone J252D4 |
| anti-human IgD | biolegend | Clone IA6-2 |
| anti-human CD27 | biolegend | Clone M-T271 |
| anti-human CD11c | biolegend | Clone S-HCL-3 |
| anti-human CD38 | biolegend | Clone HIT2 |
| anti-human CD21 | biolegend | Clone Bu32 |
| Biotin Rat Anti-Mouse CD185 (CXCR5) | BD bioscience | Clone 2G8 |
| Zombie Aqua™ Fixable Viability Kit | biolegend | Cat#423102 |
| PE Streptavidin | BD bioscience | Cat#554061 |
| T-bet (4B10) Antibody | Santa Cruz Biotechnology | Cat#sc-21749 |
| ZEB2 antibody | Novus | Cat#NBP1-82991 |

**Table S7. List of antibodies**

| Gene name | Sg sequence(5'-3') |
|---|---|
| mouse | |
| *Zeb2* Sg1 | GTACCTTCAGCGAAGCGACA |
| *Zeb2* Sg2 | TATGAATAGTAACTTGAGTG |
| *T-bet* | CGAGGACTACGCATTGCCCG |
| *Itgax* | GGGCCGTAACTCACCCTGGA |
| *Zbtb32* | CGAGGTATCGAGAGCCACAA |
| *Tfeb* | CCTCTGTGGATTACATCCGG |
| *Litaf* | GATAACAGACATACTTGCGT |
| *Nfatc2* | TTGGAGAGTGGCCACTCGAG |
| *Srebf2* | ACTCCAGTGACAGTACACTG |
| *Jazf1* | CACAGGCAGCGAGTATGATG |
| *Jun* | TGTGCCGCGGAGGTGACACT |
| *Plek* | GTTTGCCAAAGTCTTGACAA |
| *Batf* | AGAGATCAAACAGCTCACCG |
| *Tfec* | CATCAGTGGACTACATCAAG |
| *Sox5* | CGAGGGTCCGCTGGTCAGGA |
| *Mbd4* | TCAGAGTCGCCAGAAAGCAG |
| *Ets1* | TGCCTGGGGAGAGCCAGTCG |
| *Fli1* | TCACGACTGAATGTCAAGGA |
| human | |
| *ZEB2* Sg1 | TTGTAGCCCCGGTCGCAGTA |
| *ZEB2* Sg2 | GGCGCAAACAAGCCAATCCC |
| *TBX21* | AAACCGCCTGTACGTCCACC |
| *BATF* | AGGACTCTACCTGTTTGCCA |
| *FLI1* | ACTCAATCGTGAGGATTGGT |
| *SREBF2* | GCTGCATTCTGGTATATCAA |
| *ETS1* | CTTACTAATGAAGTAATCCG |
| *JUN* | TGAACCTGGCCGACCCAGTG |
| *CD19* | CTGTGCTGCAGTGCCTCAA |

**Table S8. List of sgRNA sequence**

**Supplemental Table 8. List of primers**

| | Gene name | Forward | Reverse |
|---|---|---|---|
| Mouse | | | |
| | *Hopx* | 5'- CAACTTCAACAAGGTCAACAAGCAC-3' | 5'-ACCATTTCTGCGTCTGCTCCT-3' |
| | *Zeb2* | 5'-GCAGTGAGCATCGAAGAGTACC-3' | 5'-GGCAAAAGCATCTGGAGTTCCAG-3' |
| | *Tbx21* | 5'-TGTGGATGTGGTCTTGGTGG-3' | 5'-ATTGTTGGAAGCCCCCTTGT-3' |
| | *Itgax* | 5'-TTGGCTTGTGGTCCTACTGTG-3' | 5'-GGGAAGTTCTGGCTCTGCTTG-3' |
| | *Itgam* | 5'-GTGAATATGTCCTTGGGCCTGT -3' | 5'-CGGAGCCATCAATCAAGAAGACA -3' |
| | *Zbtb32* | 5'-TCCAGATACGGTGCTCCCTTCT-3' | 5'-CCAGAGAGCTTTGGAGTGGTTC-3' |
| | *Cxcl10* | 5'-ATCATCCCTGCGAGCCTATCCT-3' | 5'-GACCTTTTTTGGCTAAACGCTTTC-3' |
| | *Rpl13a* | 5'-GGGCAGGTTCTGGTATTGGAT-3' | 5'-GGCTCGGAAATGGTAGGGG-3' |
| | *Cxcl9* | 5'-AATGCACGATGCTCCTGCA-3' | 5'-AGGTCTTTGCGGGATTTGTAGTGG-3' |
| | *Bach2* | 5'-ACAGACGAAAGATGACTTGGTG-3' | 5'-CTCTGCTGAGTAACAGCTTGG-3' |
| | *Myc* | 5'-ACGGCCTTCTCTCCTTCCTC-3' | 5'-GCCTCTTCTCCACAGACACC-3' |
| | *Nfatc2* | 5'-TCATCCAACAACAGACTGCCC-3' | 5'-GGGAGGGAGGTCCTGAAAACT-3' |
| | *Junb* | 5'-TCACGACGACTCTTACGCAG-3' | 5'-CCTTGAGACCCCGATAGGGA-3' |
| | *Jun* | 5'-CCTTCTACGACGATGCCCTC-3' | 5'-GGTTCAAGGTCATGCTCTGTTT-3' |
| | *Srebf2* | 5'-AGAAAGAGCGGTGGAGTCCTTG-3' | 5'-GAACTGCTGGAGAATGGTGAGG-3' |
| | *Foxp1* | 5'-CATGCCTCTACCAATGGACAGC-3' | 5'-GAAGTCGTCACAAACCGCCTCA-3' |
| | *Zfp36l1* | 5'-CTTCCACACCATCGGCTTTTGC-3' | 5'-CACTGGGAAACCCAGCAAAGCT-3' |
| | *Aff3* | 5'-AGATGACCTGGCTTCCTCCACT-3' | 5'-GTCCAGAGGATACAAGTTGCGAG-3' |
| | *Tfeb* | 5'-CCACCCCAGCCATCAACAC-3' | 5'-CAGACAGATACTCCCGAACCTT-3' |
| | *Klf6* | 5'-GGAAGGTTGTGAGTGGCGTTTTG-3' | 5'-AGGTGGTCAGACCTGGAGAAAC-3' |
| | *Irf1* | 5'-TCCAAGTCCAGCCGAGACACTA-3' | 5'-ACTGCTGTGGTCATCAGGTAGG-3' |
| | *Irf8* | 5'-CAATCAGGAGGTGGATGCTTCC-3' | 5'-GTTCAGAGCACAGCGTAACCTC-3' |
| | *Pou2f2* | 5'-TCCTGGAGAAGTGGCTCAACGA-3' | 5'-ATGCTGGTCCTCTTCTTGCGTC-3' |
| | *Litaf* | 5'-CAAGATGATCGTGACCCAGCTG-3' | 5'-GCAGTAGTGGTCCACATCCTGT-3' |
| | *Klf3* | 5'-CCTCTCATGGTTTCCTTGTCGG-3' | 5'-CCTCTGTGGTTCAATTCCAGGC-3' |
| | *Jazf1* | 5'-CGACATAGCAGTGGCAGCCTTA-3' | 5'-TCTCTGTGGTCCAGGACTCATC-3' |
| | *Plek* | 5'-GGAGCAGTTCACTTGAGAGGCT-3' | 5'-TGGAAGTGGCTGCCTGCAAGTA-3' |
| | *Plscr1* | 5'-GACCTCTGAGATGCAGTAGCTG-3' | 5'-GGAGAGTGAACTTGGGCAGACA-3' |
| | *Zfp36l2* | 5'-AAGTACGGCGAGAAGTGCCAGT-3' | 5'-AGAAGCCGATGGTGTGGAAGGT-3' |
| | *Fos* | 5'-AGGGGCAAAGTAGAGCAGCTA-3' | 5'-CAATCTCAGTCTGCAACGCA-3' |
| | *Tox* | 5'-AAGATGGCGCACTGCTCTCCAA-3' | 5'-CATGCTTGCCTGCTGTCTGATG-3' |
| | *Bhlhe40* | 5'-CGGATTAACGAGTGCATTGCCC-3' | 5'-GCTTCAACGTAAGCTCCAGAACC-3' |
| | *Foxp4* | 5'-CACCAGGATGTTCGCCTACTTC-3' | 5'-TATTCCCGCTCATCCACAGTCC-3' |
| | *Sox5* | 5'-CGCCAGATGAAAGAGCAACTCAG-3' | 5'-TGAGTCAGGCTCTCCAGTGTTG-3' |
| | *Crem* | 5'-GCAGCACAATCAGCCGATGGTA-3' | 5'-AGCTCGGATCTGGTAAGTTGGC-3' |
| | *Arid5a* | 5'-CGCCTCTGGAAGAACGTGTATG-3' | 5'-TGGTAGGAGGCAGTGGCTTGTC-3' |
| | *Arid3a* | 5'-TCCATCACCAGTGCTGCCTTCA-3' | 5'-TCCCTGCGATTGCTGTCTATGG-3' |
| | *Fli1* | 5'-CCATACAGACCAGTCCTCACGA-3' | 5'-CATGGTCTGTGATCCTCCAAGG-3' |
| | *Mbd4* | 5'-ACAGGATGGCTCTGAAATGCCC-3' | 5'-ACTTGTGTCCGTGGGATGCTGT-3' |
| | *Ets1* | 5'-CCAGAATCCTGTTACACCTCGG-3' | 5'-CAGCGTCTGATAGGACTCTGTG-3' |
| | *Hif1a* | 5'-CCTGCACTGAATCAAGAGGTTGC-3' | 5'-CCATCAGAAGGACTTGCTGGCT-3' |
| | *Mef2b* | 5'-AAGGTCCTGGAGAGAAGCTGCT-3' | 5'-GTAGGACAGCTCTGAAACCGAC-3' |
| | *S1pr2* | 5'-CTCACTGCTCAATCCTGTCATC-3' | 5'-TTCACATTTTCCCTTCAGACC-3' |
| | *Il21* | 5'-GATCCTGAACTTCTATCAGCTCCAC-3' | 5'-GGCATTTAGCTATGTGCTTCTGTT-3' |
| | *Ifng* | 5'-ATGAACGCTACACACTGCATC-3' | 5'-CCATCCTTTTGCCAGTTCCTC-3' |
| | *Il4* | 5'-ACTTGAGAGAGATCATCGGCA-3' | 5'-AGCTCCATGAGAACACTAGAGTT-3' |
| | *Il6* | 5'-GATGGATGCTACCAAACTGGAT-3' | 5'-CCAGGTAGCTATGGTACTCCAGA-3' |
| | *Csf2* | 5'-GTCTCTAACGAGTTCTCCTTCA-3' | 5'-TAGTAGCTGGCTGTCATGTTC-3' |
| | *Tnfa* | 5'-TCTTCTCATTCCTGCTTGTGG-3' | 5'-GGTCTGGGCCATAGAACTGA-3' |
| | *Il10* | 5'-TGAGAAGCTGAAGACCCTCA-3' | 5'-ACCTTGGTCTTGGAGCTTATT-3' |
| | *Tgfb1* | 5'-TGGAGCAACATGTGGAACTC-3' | 5'-CAGCAGCCGGTTACCAAG-3' |
| Human | | | |
| | *ZEB2* | 5'-AATGCACAGAGTGTGGCAAGGC-3' | 5'-CTGCTGATGTGCGAACTGTAGG-3' |
| | *RPL13A* | 5'-CGAGGTTGGCTGGAAGTACC-3' | 5'-CTTCTCGGCCTGTTTCCGTAG-3' |

**Table S9. List of primers**

**Reference：**


56. J. Klarquist, E. M. Janssen, The bm12 Inducible Model of Systemic Lupus Erythematosus (SLE) in C57BL/6 Mice. *J Vis Exp*, e53319 (2015).
57. A. Cossarizza *et al.*, Guidelines for the use of flow cytometry and cell sorting in immunological studies (second edition). *Eur J Immunol* **49**, 1457-1973 (2019).
58. A. P. Heinen *et al.*, Improved method to retain cytosolic reporter protein fluorescence while staining for nuclear proteins. *Cytometry A* **85**, 621-627 (2014).
59. T. Hsiau *et al.*, Inference of CRISPR edits from Sanger trace data. *BioRxiv*, (2019).
60. G. X. Zheng *et al.*, Massively parallel digital transcriptional profiling of single cells. *Nat Commun* **8**, 14049 (2017).
61. T. Stuart *et al.*, Comprehensive Integration of Single-Cell Data. *Cell* **177**, 1888-1902.e1821 (2019).
62. D. Kim, B. Langmead, S. L. Salzberg, HISAT: a fast spliced aligner with low memory requirements. *Nat Methods* **12**, 357-360 (2015).
63. S. Anders, P. T. Pyl, W. Huber, HTSeq--a Python framework to work with high-throughput sequencing data. *Bioinformatics* **31**, 166-169 (2015).
64. S. Henikoff, J. G. Henikoff, H. S. Kaya-Okur, K. Ahmad, Efficient chromatin accessibility mapping in situ by nucleosome-tethered tagmentation. *Elife* **9**, (2020).
65. M. I. Love, W. Huber, S. Anders, Moderated estimation of fold change and dispersion for RNA-seq data with DESeq2. *Genome Biol* **15**, 550 (2014).
66. H. S. Kaya-Okur *et al.*, CUT&Tag for efficient epigenomic profiling of small samples and single cells. *Nat Commun* **10**, 1930 (2019).
67. B. Langmead, S. L. Salzberg, Fast gapped-read alignment with Bowtie 2. *Nat Methods* **9**, 357-359 (2012).
68. F. Ramírez *et al.*, deepTools2: a next generation web server for deep-sequencing data analysis. *Nucleic Acids Res* **44**, W160-165 (2016).
69. Y. Zhang *et al.*, Model-based analysis of ChIP-Seq (MACS). *Genome Biol* **9**, R137 (2008).
